# Variation in infant subcortical brain development from 6 to 12 months in Down syndrome

**DOI:** 10.64898/2026.06.16.732759

**Authors:** Dea Garic, Mengwei Ren, Zoë Hawks, Yoonmi Hong, Carolyn Lasch, Rebecca Grzadzinski, Sun Hyung Kim, Omar Azrak, Jed Elison, Jason Wolff, John R. Pruett, Robert C. McKinstry, Annette Estes, Stephen Dager, Juhi Pandey, Robert T. Schultz, Alan Evans, Mark Shen, Martin Styner, Joseph Piven, Kelly Botteron, Heather Hazlett, Guido Gerig, Natasha Marrus

## Abstract

**Introduction:** Down syndrome (DS), arising from Trisomy 21, is the most common genetic condition associated with intellectual disability. While smaller total brain volumes have been consistently observed in DS, no longitudinal neuroimaging studies have examined volumetric brain development in DS during infancy, a period of rapid neural growth when interventions may have the greatest impact.

**Method:** High-resolution T1- and T2-weighted images were acquired during natural sleep in a multisite longitudinal cohort of 44 infants with DS and 39 control infants without DS at ages 6 and 12 months. Neuroimaging data were harmonized to reduce batch effects, and a novel deep-learning, repeated-measures segmentation approach was applied to optimize neuroanatomical segmentations. Total intracranial volume (ICV) and bilateral absolute subcortical volumes (amygdala, caudate, hippocampus, pallidum, putamen, thalamus) were first directly compared in infants with and without DS at 6 and 12 months. Hierarchical linear modeling (HLM) evaluated longitudinal group differences for each structure, accounting for sex, gestational age, and laterality. Subcortical group differences estimated by HLM were also compared to group differences in total ICV.

**Results:** ICV in infants with DS was lower than controls at 6 months (12.6%; *p<*.001) and 12 months (16.3%; *p<*.001). Subcortical structures displayed a range of lower volumes (6.9%-13.1%; *p’s*≤.003) in infants with DS, although the caudate and putamen were exceptions. Caudate volumes were on average lower in DS but not significantly different from controls, while putamen volumes were on average higher in DS but not significantly different from controls, except for the right putamen, which was significantly larger (5.3%; *p*=.018) at 6 months. In HLM, ICV and all subcortical structures showed slower growth in DS from 6 to 12 months, except for the amygdala and putamen, which displayed similar growth rates to controls. DS-associated reductions in subcortical volumes were similar in magnitude to ICV, although 12-month caudate and 6- and 12-month putamen volumes were enlarged relative to ICV.

**Conclusion:** Infants with DS exhibited substantially reduced ICV and widespread reductions in subcortical volumes and growth from 6-12 months. Across a range of volumetric differences, findings were most distinct in the basal ganglia, for which volume reductions were attenuated in the caudate, while the putamen was uniquely enlarged with comparable growth to controls. These observations support early regional specificity in the neural impact of Trisomy 21 and underscore the utility of infant neuroimaging to inform biologically based interventions and clinical trial readiness in DS.

## 1. INTRODUCTION

Down syndrome (DS), which arises from a triplication of chromosome 21, is the most common genetic condition leading to intellectual disability (ID) worldwide, with a prevalence rate of 1 in 700 live births.^1,2^ While ID is a highly penetrant phenotype in DS, there is growing recognition of heterogeneity and a distinct DS-associated phenotypic profile that includes relative weaknesses in verbal processing,^3,4^ working memory,^4–7^ and attention,^8^ and relative strengths in nonverbal social functioning^9–11^ and visuospatial processing.^12,13^ Behavioral delays preceding ID are present during infancy, a period of accelerated brain and behavioral development, and are followed by the long-term impact of ID on function and quality of life.^3,6,8,14,15^ Beyond early cognitive impairment from ID, approximately 50% of individuals with DS later develop Alzheimer’s dementia decades earlier than the general population,^16–19^ which is attributed to a 1.5-fold excess in amyloid beta production caused by an extra copy of the amyloid precursor protein gene on chromosome 21.^20,21^ Accumulating progress in animal studies^22–25^ suggests actionable opportunities via genetic targets on chromosome 21 for biologically based treatment of cognitive impairments in DS. Characterizing the early foundations of brain differences due to Trisomy 21 (T21) is crucial for clarifying the neural underpinnings of cognitive impairments and guiding interventions oriented to specific needs across the lifespan.

Despite cognitive delays by infancy, there is a lack of longitudinal neuroimaging studies in DS during this period, when rapid increases in gray matter volumes, myelination and synaptogenesis,^26,27^ along with heightened neuroplasticity, could allow interventions to have the largest impact. Neuropathology studies of fetal brain development in DS have revealed significant disruptions, including atypical neural differentiation and reduced neurogenesis, myelination, dendritogenesis, and synaptogenesis in cortical and subcortical brain regions.^28–31^ Developmental neuroimaging thus provides an important opportunity to extend early neural characterization to structural brain development, trajectories, and associations with behavioral outcomes. However, the few infant and early childhood neuroimaging studies in DS have been constrained by cross-sectional methodologies, small samples (*n’*s<20), and reliance on clinically obtained neuroimaging, which can increase bias and reduce standardization of data collection.^32^ These limitations have hindered the discovery of neural biomarkers that could predict a range of later neurodevelopmental outcomes for young children with DS, as well as help disentangle neurodevelopmental vs. neurodegenerative brain differences relevant to patient selection in future clinical trials.

Most neuroimaging studies characterizing brain phenotypes in DS entail volumetric studies. The most consistent DS-associated brain finding is substantially smaller overall brain volume than the general population, which has been observed as early as the second trimester in utero,^33–36^ as well as in newborns,^37^ children,^32,38,39^ and adults.^40–42^ A recent study^42^ that divided participants with DS into age groups from early childhood through adulthood found moderate-to-large effect sizes for smaller volumes, with more modest effects in early childhood (Cohen’s *d*=-0.65), and the highest effects in adults (Cohen’s *d*=-1.35). These observations suggest early emergence of brain volumetric differences followed by a widening gap between childhood and adulthood, implying slower brain growth in DS.

Compared with overall brain volume, characterization of subcortical volumes of structures, including the amygdala, hippocampus, thalamus, and basal ganglia, has been limited in DS. Only one prior cross-sectional study has examined subcortical volumes in infants with DS between 32–46 weeks postmenstrual age.^37^ These structures, each with their respective embryologic origins and functions, warrant developmental examination in DS, given their associations with outcomes in neurodevelopmental disorders^43–47^ as well as functional domains centrally affected in DS, including motor development,^48–52^ emotion regulation,^53,54^ verbal processing,^55,56^ and working memory.^57,58^ Further, these structures undergo rapid, regionally specific growth in infancy,^52,53^ which could inform developmentally sensitive neural biomarkers and guide future DS intervention targets.

Among subcortical structures, relatively consistent DS-related differences have been reported in the hippocampus, lentiform nucleus, and caudate. Multiple studies have reported hippocampal volume reductions in children with DS compared with controls,^41,59–61^ whereas basal ganglia volumes appear less likely to show DS-associated reductions. The lentiform nucleus, composed of the pallidum and putamen, appears to be consistently larger throughout development in infants and children with DS than in controls,^37,41,42,60^ with putamen enlargement observed into adulthood.^42,62^ Findings for the caudate in DS show volumes similar to controls^37,41^ or relative increases when considering overall reduced brain size.^42^

In contrast, mixed volumetric findings have been observed for the amygdala and thalamus, with studies reporting relatively larger^37,42^ or smaller^38,41,59,63,64^ volumes, or no difference^37,41,42,61^ compared to controls. In addition to methodological variability in neuroimaging sequences, structural segmentations, and anatomical definitions, small sample sizes and age ranges spanning childhood to adulthood^41,61,64–70^ may contribute to these discrepancies. Altogether, prior work supports regional variation in DS-associated differences in cortical and subcortical volumes,^41,66^ and a potential dissociation between relatively typical volumes in the basal ganglia versus whole brain and some subcortical regions. However, larger samples, developmentally defined age ranges, rigorous segmentations, and longitudinal designs are needed to clarify the extent to which these features manifest by infancy and whether relative growth rates in infants with DS align with volumetric differences observed at later ages.

Here we report on what is, to our knowledge, the first longitudinal study to characterize differences in subcortical brain volume and development between infants with and without DS, at 6 and 12 months, a period of rapid volumetric expansion.^53^ High-resolution neuroimaging data, acquired with identical scanners and imaging protocols, were collected in infants with DS and controls during natural sleep across four study sites, and a novel deep-learning-based, repeated-measures segmentation was used to maximize the quality of infant anatomical structure segmentations.^71^ We examined (1) 6- and 12-month group differences in total intracranial volume (ICV) and bilateral absolute subcortical volumes across amygdala, caudate, hippocampus, pallidum, putamen, and thalamus, (2) group differences in 6- to 12-month longitudinal change in ICV and these subcortical volumes and (3) subcortical volumetric group differences relative to group differences in total ICV to distinguish global from regionally specific effects of DS. Given prior literature, we hypothesized smaller ICV in DS than in controls, regional variation in group differences in subcortical volumes, and slower rates of growth in ICV and subcortical volumes in DS. Characterization of global and subcortical differences during infancy in DS provides critical context for future neurobiologically based treatments and sensitive timeframes for intervention.

## 2. Method

### 2.1 Participants

This study was conducted as part of the Infant Brain Imaging Study of Down syndrome (IBIS-DS), an ongoing multi-site longitudinal study to characterize infant brain and behavioral development in Down syndrome relative to a typically developing control group. Data collection took place across four sites: Washington University in St. Louis (WUSTL), University of North Carolina at Chapel Hill (UNC), Children’s Hospital of Philadelphia (CHOP), and University of Washington (UW). McGill University served as the Data Coordinating Center and utilized the LORIS data management platform^72^ as the hub for multimodal data collection, curation, preparation, and archiving.

Infants with DS were reported by parents to have a diagnosis of full T21 (i.e., not partial or mosaic T21). Infants in the control sample did not have DS or any first-degree relatives with a history of autism or intellectual disability. Participants in the control sample were recruited concurrently. Exclusion criteria for both the DS and control groups included: (1) a diagnosis or physical signs of known genetic conditions (other than DS in the DS group), (2) significant medical conditions that could impact growth, development, cognition, or sensory function (except for conditions commonly associated with DS in DS infants), (3) birth weight <2500 grams, gestational age <34 weeks for DS infants or <36 weeks for control infants, (4) history of significant perinatal complications, (5) prenatal exposure to neurotoxins, (6) maternal gestational diabetes requiring medication, (7) contraindication for MRI, and (8) families whose primary language is not English, due to requirements for cognitive assessments.

Ethical approval was obtained from institutional review boards at all sites, which relied on the single IRB at Washington University in St. Louis. Written informed consent was obtained from at least one parent.

### 2.2 Neuroimaging Protocols

All imaging data were acquired in 6- and 12-month-old infants on Siemens 3.0T Prisma MRI scanners at all sites using a 32-channel head coil. T1-weighted MP-RAGE (TR = 2400ms, TI = 1060ms, TE = 2.24ms, 0.8mm^3^ voxel, TA: 6:38) and T2-weighted 3DTSE SPACE (TR = 3200ms, TE = 564ms, 0.8mm^3^ voxel, TA: 5:57) images were used in volumetric analyses. Infants were imaged during natural sleep without sedation and wore pediatric earplugs in addition to headphones for hearing protection. Background MRI sounds were played over the scan room speakers to minimize infants’ startle responses to the sudden onset of scanner noise. Extra care was taken with the head position (e.g., scanning infants on an incline) to optimize breathing and minimize snoring, which is common in DS and can introduce movement artifacts in imaging data.

#### Qualitative MRI/Radiological Review

MR images were reviewed for clinically relevant abnormalities and incidental findings by a board-certified neuroradiologist at each site.

#### Structural MRI Processing

Preprocessing included intensity inhomogeneity correction, pediatric ICBM space alignment, and skull stripping. Skull stripping was performed via an automated convolutional neural network in ANTsPyNet^73,74^ followed by manual correction where necessary.

#### Temporally-specific segmentation

We applied a novel, deep-learning-based, temporally-specific segmentation method^71^ that embeds longitudinal processing into the image segmentation step and incorporates the self-similarity of structures of individual subjects repeatedly imaged at different time points. This deep-learning approach has been shown to significantly improve the segmentation quality of anatomical structures,^71^ consistent with previous work demonstrating that accounting for non-independence in longitudinal segmentation data can increase effect sizes in statistical analyses.^75^ Multiple key innovations to improve consistency of structural segmentations for individuals across multiple time points included: a) presentation of a novel spatiotemporal representation learning procedure that exploits the spatiotemporal self-similarity of multi-scale intra-subject image features, b) integration of self-supervised consistency regularization to utilize intra-subject anatomical correlation, and c) learning from few annotations to eliminate the necessity to train age-specific atlases with large annotated datasets, a long-recognized obstacle for processing of infant image data that features rapidly changing contrast and shape across age. The approach is robust to batch effects and site variance, resulting in only limited residual site effects that required further harmonization (see Supplementary Material, Figure 1). The network was trained on legacy IBIS training datasets (386 subjects with 887 images from subjects imaged at 6, 12, and 24 months) following work by Hazlett and colleagues.^76^ Processing of the total of 232 T1w/T2w MRI datasets of DS and control subjects in this study was blind to group and/or diagnosis.

#### Data harmonization

While identical scanners and imaging protocols were used at every site, as well as a segmentation method largely robust to batch and site effects, residual site differences can still potentially introduce non-biological variability in the data. Therefore, all data extracted from the images in this study were harmonized across sites using neuroComBat,^77^ which has been shown to effectively reduce site-related variance while preserving biological variability.^78^ Because neuroCombat removes site-specific effects, site was not included as a covariate in subsequent statistical models.

#### Data Analysis

##### Descriptive analyses of participant characteristics and cross-sectional brain development

Sociodemographic characteristics, as well as ICV and absolute bilateral subcortical volumes for amygdala, caudate, hippocampus, pallidum, putamen, and thalamus were directly compared in infants with and without DS. Continuous variables were compared using independent samples t-tests; χ^2^ tests were used for categorical variables. Cohen’s *d* effect sizes were calculated to index group differences in volumes for infants with and without DS at 6 and 12 months. At each time point, *p*-values were adjusted for multiple comparisons using false discovery rate correction^79^ and reported as *q*-values, with 13 comparisons each at 6 and 12 months (ICV plus bilateral volumes for 6 subcortical structures).

##### Longitudinal analyses of brain development

We applied hierarchical linear modeling (HLM), a regression technique that accounts for shared variance in nested data,^80^ to test for differences in the rate of 6- to 12-month volumetric change between infants with and without DS. Neuroimaging observations at 6 and 12 months were nested within individuals. Analyses were performed in R v4.3.1,^81^ using the tidyverse package for data wrangling and visualization^82^ and the lme4 package for HLM.^83^

We first examined relationships for ICV, including cohort (control, DS) as a moderator of the association between ICV (z-scored) and age in months. Models included a random intercept and covaried for sex and age at birth. To preserve statistical power, median imputation by cohort was conducted for missing gestational ages in eight participants.

Separate analogous models were applied to each subcortical region and included hemisphere (left, right) and cohort as moderators of the association between gray matter volume (z-scored) and age in months. Absolute volumes, rather than ICV-adjusted volumes, were used to examine the impact of DS on specific structures, given evidence for non-uniform reductions in cellularity across brain regions^84^ in DS, which could complicate interpretability. All models (i.e., ICV and subcortical volumes) tested two-way interactions between age and cohort. Subcortical models additionally tested three-way interactions with laterality (left and right hemispheres [LH, RH]). To account for multiple comparisons, *p*-values were adjusted for seven models (ICV plus 6 subcortical structures) using false discovery rate correction,^79^ and the adjusted p-values (*q*-values) were reported.

In addition to evaluating longitudinal changes in volume via age-by-cohort interactions, we ran models under different centering conditions for age (6 and 12 months) and with different hemisphere reference levels to determine group differences at each age for bilateral subcortical volumes.

##### Comparison of group differences in subcortical volumes relative to ICV

In accordance with field standards,^37^ group differences in subcortical volumes were also examined relative to those in ICV. Covariate-adjusted absolute values from the HLM were used to qualitatively compare the standardized main effect of cohort for each subcortical region to that of ICV, indicating whether DS-associated regional differences were comparable to, larger than, or smaller than overall ICV differences. This was applied as the primary approach versus covarying for ICV within HLM because relationships between subcortical volumes and ICV can differ with time or by diagnostic group, potentially leading to misinterpretation of age interactions within linear models. As a supplemental analysis, models with ICV as a covariate were run to provide quantitative group-level comparisons for each time point.

## 3. Results

### 3.1 Participant Characteristics

Infants with DS (n=44) and control infants without DS (n=39) with brain volumetric data at 6 and/or 12 months of age (Table 1) did not differ by sex (*p*=.10), age at scan at either time point (*p’*s=.06 and .75 for 6 and 12 months, respectively), maternal education (*p*=.33), race (*p*=.96), ethnicity (*p*=.44), or household income (*p*=.16). Mean gestational age was significantly lower in the DS versus control sample (38 weeks vs. 39 weeks, *p*<.001) and was included as a covariate in all models.

**Table 1:** Participant Demographics by Diagnostic Outcome Group The groups did not differ on sex (*p* = 0.10), age at scan for either time point (*ps* = 0.06 and 0.75 for 6 and 12 months, respectively), maternal education (*p* = 0.33), race (*p* = 0.96), ethnicity (*p* = 0.44), nor household income (*p* = 0.16). Gestational age was significantly lower in the DS sample (38 weeks compared to 39 weeks, *p* < 0.001) and was used as a covariate in all models. Age and sex data are represented a mean and standard deviation (SD) in parentheses. All other data is presented as *n* and the percentage of the group. F, female; M, male; MRI, magnetic resonance imaging.

|  | **Down Syndrome** | **Control** | **Test Statistic or Total** |
|---|---|---|---|
| ***n* at 6-12 month MRI visits** | 44 | 39 | **Total = 83** |
| *Sex (M/F)* | 17/27 | 22/17 | ***χ*<sup>2</sup> = 2.62, *p* = .10** |
| *Gestational age weeks, M (SD)* | 38.08 (1.39) | 39.07 (1.20) | ***T* = 3.45, *p* = .0009** |
| *Age at 6-month MRI, M (SD)* | 6.86 (1.07) | 6.50 (.47) | ***T* = 1.88, *p* = .06** |
| *Age at 12-month MRI, M (SD)* | 12.67 (1.24) | 12.57 (.64) | ***T* = 0.32, *p* = .75** |
| **Maternal Education; M (SD)** | 4.40 (1.40) | 4.68 (1.14) | ***T* = 0.98, *p* = .33** |
| *1) Some high school* | *0* | *0* |  |
| *2) High school graduate* | *4 (9%)* | *0* |  |
| *3) Some college* | *8 (18%)* | *5 (13%)* |  |
| *4) College graduate* | *14 (32%)* | *17 (44%)* |  |
| *5) Some grad school* | *1 (2%)* | *1 (3%)* |  |
| *6) Graduate degree* | *16 (36%)* | *15 (38%)* |  |
| *NA) Not answered* | *1 (2%)* | *1 (3%)* |  |
| **Race** |  |  | **Fisher’s, *p* =.96** |
| *Asian* | *2 (4%)* | *2 (5%)* |  |
| *Black/African American* | *3 (7%)* | *4 (10%)* |  |
| *White* | *34 (77%)* | *29 (74%)* |  |
| *More than one race* | *3 (7%)* | *3 (8%)* |  |
| *Not answered* | *2 (4%)* | *1 (3%)* |  |
| **Ethnicity** |  |  | **Fisher’s, *p* =.44** |
| *Hispanic* | *5 (11%)* | *2 (5%)* |  |
| *Non-Hispanic* | *37 (84%)* | *35 (90%)* |  |
| *Not answered* | *2 (4%)* | *2 (5%)* |  |
| **Household Income:** | 5.61 (1.95) | 6.16 (1.40) | ***T* = 1.42, *p* = .16** |
| *1) Less than 25k* | *3 (7%)* | *0* |  |
| *2) 25k - 35k* | *0* | *0* |  |
| *3) 35k - 50k* | *2 (4%)* | *0* |  |
| *4) 50k – 75k* | *6 (14%)* | *6 (15%)* |  |
| *5) 75k – 100k* | *5 (11%)* | *6 (15%)* |  |
| *6) 100k – 150k* | *10 (23%)* | *10 (26%)* |  |
| *7) 150k – 200k* | *8 (18%)* | *6 (15%)* |  |
| *8) Over 200k* | *7 (16%)* | *9 (23%)* |  |
| *NA) Not answered* | *3 (7%)* | *2 (5%)* |  |

F, female; M, male; MRI, magnetic resonance imaging.

### 3.2 Group Comparisons of Absolute Volumes at 6 and 12 months

Descriptive statistics for total ICV and absolute bilateral subcortical volumes by DS and control groups are provided in **Table 2**. Total ICV was significantly lower in infants with DS at 6 months (12.6% decrease) and 12 months (16.3% decrease). These differences comprised large effect sizes of *d* = -1.47 and *d* = -2.03, respectively (*q’s*<.001).

**Table 2.** 6- and 12-month gray matter volumes (mm3) in control and Down syndrome (DS) cohorts. Volumes formatted as: mean (standard deviation), minimum – maximum. Ratios are calculated based on absolute volume relative to total intracranial volume (ICV). Values below 100% indicate lower DS volumes than in controls. Bolded values passed multiple comparison correction. ^a^FDR corrected p<.001, ^b^FDR corrected p<.01, ^c^FDR corrected p<.05

|  | 6 Months |  |  |  | 12 Months |  |  |  |
|---|---|---|---|---|---|---|---|---|
|  | Control (n=37) | DS (n=39) | % Relative Volume (DS vs. Control) | Statistics | Control (n=20) | DS (n=20) | % Relative Volume (DS vs. Control) | Statistics |
| ICV | 913348.9 (79198.8)<br>766000.1 - 1086799.9 | 798024.6 (76471.1)<br>628908.7 - 941746.7 | 87.4% | ***t*(74)=-6.46; *p*<.001<sup>a</sup>; *d*=-1.47** | 1082217.3 (97492.6)<br>900325.1 - 1271916.1 | 905759.0 (70641.4)<br>790533.1 - 1044680.2 | 83.7% | ***t*(38)=-6.56; *p*<.001<sup>a</sup>; *d*=-2.03** |
| Left Amygdala | 789.0 (58.9)<br>641.3 - 936.6 | 672.52 (84.7)<br>494.3 - 871.4 | 85.2% | ***t*(74)=-6.99; *p*<.001<sup>a</sup>; *d*=-1.57** | 899.4 (70.24)<br>774.8 - 1043.2 | 782.1 (76.7)<br>680.7 - 937.8 | 87.0% | ***t*(38)=-5.04; *p*<.001<sup>a</sup>; *d*=-1.56** |
| Right Amygdala | 773.3 (60.7)<br>624.9 - 918.5 | 688.0 (81.0)<br>480.9 - 863.1 | 89.0% | ***t*(74)=-5.18; *p*<.001<sup>a</sup>; *d*=-1.47** | 898.5 (75.0)<br>796.7 - 1064.2 | 810.3 (69.3)<br>724.0 - 1006.9 | 90.2% | ***t*(38)=-3.86; *p*<.001<sup>a</sup>; *d*=-1.20** |
| Left Caudate | 2164.9 (224.0)<br>1722.0 - 2679.9 | 2111.9 (221.2)<br>1652.5 - 2747.9 | 97.6% | *t*(74)=-1.04; *p*=.303; *d*=-0.24 | 2622.8 (262.5)<br>2148.6 - 3264.3 | 2540.4 (266.8)<br>2102.8 - 3050.1 | 96.9% | *t*(38)=-0.99; *p*=.33; *d*=-0.31 |
| Right Caudate | 2246.7 (226.6)<br>1757.2 - 2781.0 | 2153.8 (234.1)<br>1666.6 - 2931.1 | 95.9% | *t*(74)=-1.76; *p*=.083; *d*=-0.40 | 2669.7 (278.1)<br>2099.6 - 3308.0 | 2580.8 (275.8)<br>2126.4 - 3117.2 | 96.7% | *t*(38)=-1.02; *p*=.32; *d*=-0.32 |
| Left Hippocampus | 1828.2 (172.0)<br>1449.8 - 2112.4 | 1634.1 (148.7)<br>1239.5 - 1983.6 | 89.4% | ***t*(74)=-5.27; *p*<.001<sup>a</sup>; *d*=-1.20** | 2028.2 (226.0)<br>1691.1 - 2464.4 | 1760.1 (131.1)<br>1570.1 - 2014.8 | 86.8% | ***t*(38)=-4.59; *p*<.001<sup>a</sup>; *d*=-1.42** |
| Right Hippocampus | 1830.4 (178.2)<br>1529.1 - 2290.1 | 1649.6 (150.4)<br>1287.3 - 1934.8 | 90.1% | ***t*(74)=-4.79; *p*<.001<sup>a</sup>; *d*=-1.09** | 2090.3 (224.2)<br>1785.3 - 2554.9 | 1824.2 (131.6)<br>1597.0 - 2062.9 | 87.3% | ***t*(38)=-4.58; *p*<.001<sup>a</sup>; *d*=-1.42** |
| Left Pallidum | 709.6 (59.4)<br>581.4 - 810.9 | 660.6 (60.8)<br>542.7 - 789.2 | 93.1% | ***t*(74)=-3.55; *p*<.001<sup>a</sup>; *d*=-0.81** | 880.6 (92.8)<br>717.2 - 1058.4 | 793.2 (64.0)<br>678.7 - 889.0 | 90.1% | ***t*(38)=-3.47; *p*=.001<sup>b</sup>; *d*=-1.08** |
| Right Pallidum | 724.9 (62.0)<br>616.4 - 851.1 | 659.9 (60.5)<br>538 - 811.4 | 91.0% | ***t*(74)=-4.63; *p*<.001<sup>a</sup>; *d*=-1.05** | 897.4 (86.8)<br>743.8 - 1037.8 | 815.2 (75.4)<br>676.2 - 938.9 | 90.8% | ***t*(38)=-3.20; *p*=.003<sup>b</sup>; *d*=-0.99** |
| Left Putamen | 2705.0 (259.3)<br>2104.2 - 3251.9 | 2793.7 (244.3)<br>2274.5 - 3351.9 | 103.3% | *t*(74)=1.53; *p*=.13; *d*=0.35 | 3343.7 (332.4)<br>2698.7 - 3976.2 | 3425.9 (257.7)<br>2834.0 - 3882.5 | 102.5% | *t*(38)=0.88; *p*=.39; *d*=0.27 |
| Right Putamen | 2742.5 (266.5)<br>2073.1 - 3182.6 | 2886.5 (254.6)<br>2458.8 - 3561.0 | 105.3% | ***t*(74)=2.41; *p*=.018<sup>c</sup>; *d*=0.55** | 3420.5 (327.2)<br>2728.9 - 3994.9 | 3472.1 (222.3)<br>2934.2 - 3887.5 | 101.5% | *t*(38)=0.58; *p*=.56; *d*=0.18 |
| Left Thalamus | 5180.7 (271.1)<br>4593.6 - 5632.6 | 4567.5 (320.7)<br>3731.8 - 5098.0 | 88.2% | ***t*(74)=-8.98; *p*<.001<sup>a</sup>; *d*=-2.04** | 5533.0 (357.6)<br>4981.8 - 6232.3 | 4808.1 (269.2)<br>4326.2 - 5526.4 | 86.9% | ***t*(38)=-7.24; *p*<.001<sup>a</sup>; *d*=-2.25** |
| Right Thalamus | 5116.2 (285.1)<br>4447.9 - 5631.8 | 4503.2 (321.9)<br>3717.8 - 5082.6 | 88.0% | ***t*(74)=-8.77; *p*<.001<sup>a</sup>; *d*=-1.99** | 5439.4 (356.6)<br>4925.5 - 6110.9 | 4728.7 (297.9)<br>4237.1 - 5516.5 | 86.9% | ***t*(38)=-6.84; *p*<.001<sup>a</sup>; *d*=-2.12** |

At both ages, all subcortical volumes, except the putamen, were numerically lower in infants with DS than in controls. These differences were significant, including when accounting for multiple comparisons, for all subcortical structures except the left and right caudate at 6 and 12 months. At 6 months, the largest reduction was observed in the left amygdala (14.8%, *d=*-1.57), the only reduction to numerically exceed total ICV. At 12 months, left and right thalamus showed the largest reduction (13.1%, *d*=-2.12 -2.25). Nominally higher absolute volumes were observed for the putamen, ranging between 3.3-5.3% at 6 months and 1.5-2.5% at 12 months, although only enlargement for the 6-month right putamen was statistically significant. **Figure 1** provides a color-scaled visualization of variability in 6- and 12-month volumetric differences for subcortical volumes in infants with versus without DS. Across timepoints, most structures showed congruent levels of group differences, although warmer colors in the hippocampus and thalamus highlighted relative volume reductions for infants with DS from 6 to 12 months.

**Figure 1.**
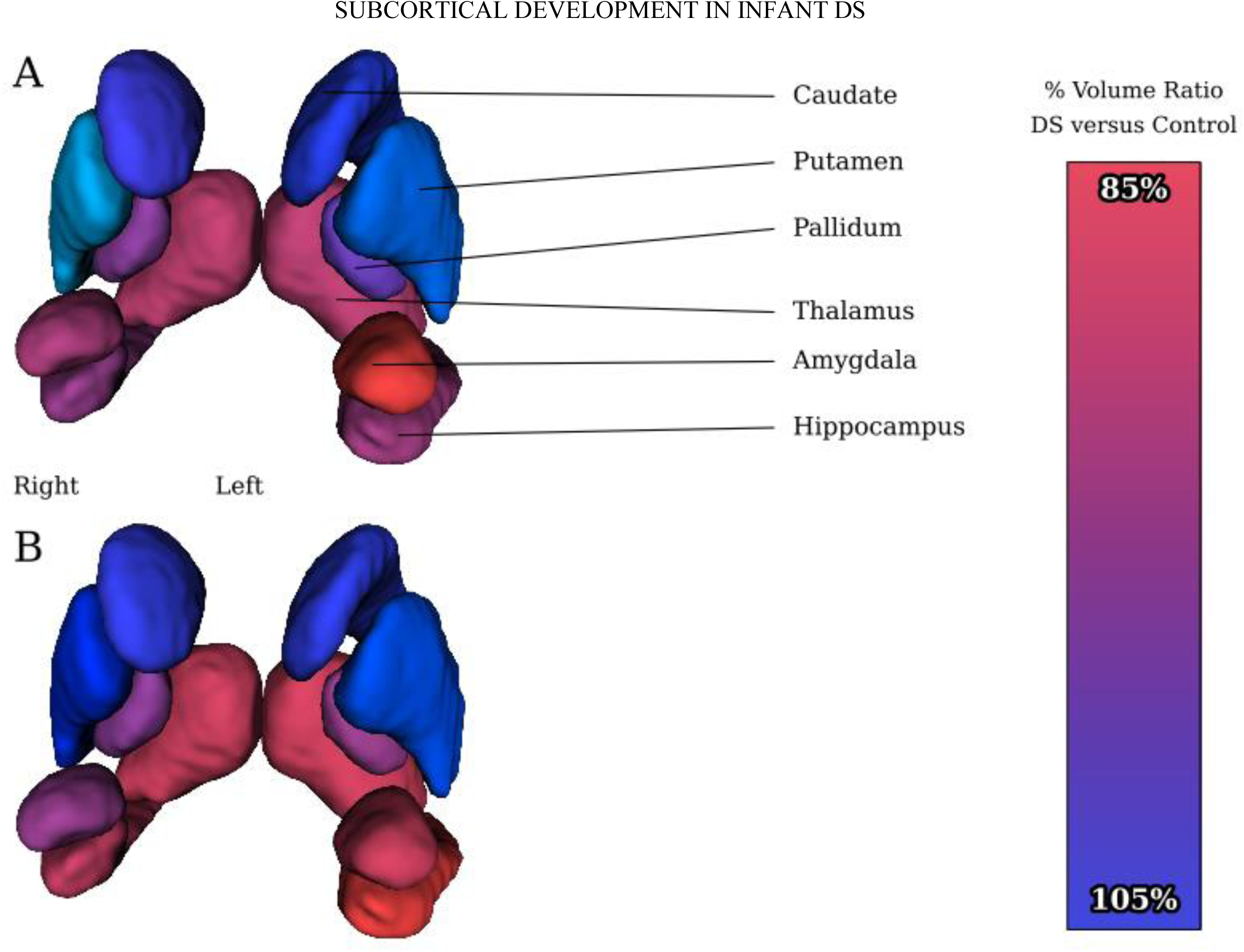
3D visualizations of subcortical volume ratios between Down syndrome (DS) and controls at 6 (A) and 12 months of age (B). A color bar on the right marks the percent range. Ratios are calculated from the absolute volumes. Values below 100% indicate lower volumes in infants with DS vs controls, with red indicating 85% of the volume observed in controls and green indicating volumes 5% larger than those in controls. Volumetric measurements and percent volume ratios per region are listed in Table 2.

### 3.3. Hierarchical Linear Modeling (HLM) and Longitudinal Group Differences

Model statistics, including those adjusted for multiple comparisons, are provided in **Table 3**, with age centered at 6 months. Sex, gestational age, and age at scan showed associations with almost all brain regions. Generally, males displayed larger volumes. Gestational age and age at scan were positively correlated with brain volume, consistent with prior literature.^27,85^

**Table 3.** Estimates (Est), standard errors (SE), and p-values (*p*) from hierarchical linear models characterizing early (6-to-12-month) gray matter development in controls and DS. Gray matter development was examined at the whole-brain level (ICV) and in six subcortical regions (amygdala, caudate, hippocampus, pallidum, putamen, and thalamus). The right hemisphere was used as a reference, centered at the 6-month time point. See **Supplementary** Tables 1-3 for left hemisphere reference and 12-month time point centering. *ICV = intracranial volume, DS = Down syndrome, Hemi = hemisphere*. ^a^FDR corrected p<.001, ^b^FDR corrected p<.01, ^c^FDR corrected p<.05

|  | ICV |  |  | Amygdala |  |  | Caudate |  |  | Hippocampus |  |  | Pallidum |  |  | Putamen |  |  | Thalamus |  |  |
|---|---|---|---|---|---|---|---|---|---|---|---|---|---|---|---|---|---|---|---|---|---|
| *Predictors* | *Est* | *SE* | *p* | *Est* | *SE* | *p* | *Est* | *SE* | *p* | *Est* | *SE* | *p* | *Est* | *SE* | *p* | *Est* | *SE* | *p* | *Est* | *SE* | *p* |
| (Intercept) | -0.22 | 0.12 | 0.076 | -0.06 | 0.15 | 0.689 | -0.60 | 0.17 | **<0.001** | -0.08 | 0.15 | 0.619 | -0.53 | 0.13 | **<0.001** | -0.91 | 0.13 | **<0.001** | 0.29 | 0.13 | **0.027** |
| Sex [Male] | 0.59 | 0.12 | **<0.001<sup>b</sup>** | 0.43 | 0.14 | **0.003<sup>b</sup>** | 0.09 | 0.17 | 0.612 | 0.48 | 0.14 | **0.001<sup>b</sup>** | 0.31 | 0.13 | **0.014<sup>c</sup>** | 0.35 | 0.13 | **0.006<sup>b</sup>** | 0.51 | 0.13 | **<0.001<sup>b</sup>** |
| Gestational age | 0.14 | 0.05 | **0.003<sup>c</sup>** | 0.10 | 0.05 | 0.076 | 0.01 | 0.06 | 0.925 | 0.14 | 0.05 | **0.013<sup>c</sup>** | 0.07 | 0.05 | 0.177 | 0.11 | 0.05 | **0.018<sup>c</sup>** | 0.13 | 0.05 | **0.008<sup>c</sup>** |
| Cohort [DS vs Control] | -0.66 | 0.14 | **<0.001<sup>b</sup>** | -0.92 | 0.17 | **<0.001<sup>a</sup>** | -0.17 | 0.19 | 0.366 | -0.62 | 0.17 | **0.001<sup>b</sup>** | -0.36 | 0.14 | **0.012<sup>c</sup>** | 0.29 | 0.14 | **0.040<sup>c</sup>** | -1.04 | 0.14 | **<0.001<sup>a</sup>** |
| age (6mo) | 0.23 | 0.01 | **<0.001<sup>a</sup>** | 0.19 | 0.02 | **<0.001<sup>a</sup>** | 0.25 | 0.01 | **<0.001<sup>a</sup>** | 0.16 | 0.02 | **<0.001<sup>a</sup>** | 0.26 | 0.01 | **<0.001<sup>a</sup>** | 0.25 | 0.01 | **<0.001<sup>a</sup>** | 0.15 | 0.01 | **<0.001<sup>a</sup>** |
| Cohort [DS] × age (6mo) | -0.08 | 0.02 | **<0.001<sup>b</sup>** | -0.03 | 0.02 | 0.170 | -0.04 | 0.01 | **0.003<sup>b</sup>** | -0.08 | 0.03 | **0.006<sup>b</sup>** | -0.05 | 0.02 | **0.002<sup>b</sup>** | 0.02 | 0.01 | 0.142 | -0.06 | 0.01 | **<0.001<sup>b</sup>** |
| hemi [R vs L] |  |  |  | -0.17 | 0.08 | **0.029<sup>c</sup>** | 0.27 | 0.05 | **<0.001<sup>b</sup>** | -0.01 | 0.10 | 0.904 | 0.13 | 0.06 | **0.018<sup>c</sup>** | 0.09 | 0.05 | 0.062 | -0.14 | 0.05 | **0.003<sup>b</sup>** |
| Cohort [DS] × hemi [R] |  |  |  | 0.28 | 0.11 | **0.012<sup>c</sup>** | -0.14 | 0.07 | **0.035** | 0.03 | 0.14 | 0.800 | -0.19 | 0.08 | **0.019<sup>c</sup>** | 0.16 | 0.06 | **0.017<sup>c</sup>** | 0.01 | 0.07 | 0.833 |
| age (6mo) × hemi [R] |  |  |  | 0.03 | 0.02 | 0.168 | -0.02 | 0.01 | 0.120 | 0.04 | 0.02 | 0.070 | 0.01 | 0.01 | 0.714 | 0.02 | 0.01 | 0.175 | -0.01 | 0.01 | 0.451 |
| (Cohort [DS] × age (6mo) × hemi [R] |  |  |  | 0.00 | 0.03 | 0.984 | 0.02 | 0.02 | 0.224 | -0.00 | 0.03 | 0.963 | 0.04 | 0.02 | 0.062 | -0.03 | 0.02 | **0.037** | -0.00 | 0.02 | 0.989 |
| **Random Effects** |  |  |  |  |  |  |  |  |  |  |  |  |  |  |  |  |  |  |  |  |  |
| σ<sup>2</sup> | 0.06 |  |  | 0.10 |  |  | 0.03 |  |  | 0.15 |  |  | 0.05 |  |  | 0.03 |  |  | 0.03 |  |  |
| τ<sub>00</sub> | 0.24<sub>IDs</sub> |  |  | 0.36<sub>IDs</sub> |  |  | 0.56<sub>IDs</sub> |  |  | 0.35<sub>IDs</sub> |  |  | 0.30<sub>IDs</sub> |  |  | 0.30<sub>IDs</sub> |  |  | 0.32<sub>IDs</sub> |  |  |
| ICC | 0.81 |  |  | 0.79 |  |  | 0.94 |  |  | 0.70 |  |  | 0.86 |  |  | 0.90 |  |  | 0.91 |  |  |
| N | 83<sub>IDs</sub> |  |  | 83<sub>IDs</sub> |  |  | 83<sub>IDs</sub> |  |  | 83<sub>IDs</sub> |  |  | 83<sub>IDs</sub> |  |  | 83<sub>IDs</sub> |  |  | 83<sub>IDs</sub> |  |  |
| Observations | 116 |  |  | 232 |  |  | 232 |  |  | 232 |  |  | 232 |  |  | 232 |  |  | 232 |  |  |
| Marginal R<sup>2</sup> / Conditional R<sup>2</sup> | 0.720 / 0.947 |  |  | 0.584 / 0.912 |  |  | 0.444 / 0.967 |  |  | 0.526 / 0.860 |  |  | 0.656 / 0.951 |  |  | 0.668 / 0.967 |  |  | 0.674 / 0.969 |  |  |

Consistent with group comparisons of absolute volumes, HLM showed significant DS-associated reductions in ICV and most subcortical volumes. The two exceptions were the caudate, which showed a non-significant negative relationship, and the putamen, which showed a significant positive relationship with DS, corresponding to larger putamen volumes in infants with DS.

Significant age-by-cohort interactions indicated slower 6- to 12-month growth in several volumes in infants with DS compared to controls. Decreased volumetric growth in DS was observed in ICV and all subcortical structures except the amygdala and putamen, for which growth rates did not differ from controls (**Figure 2**). Compared to controls, infants with DS also exhibited altered laterality in the amygdala, pallidum, putamen, and caudate (though this region did not survive FDR-correction, *q*=.052) (**Table 3**). DS was associated with right lateralization of the putamen, lower right lateralization of the caudate and pallidum, and right as opposed to left lateralization observed in infants without DS for the amygdala (**Supplemental Tables 1-3**).

**Figure 2.**
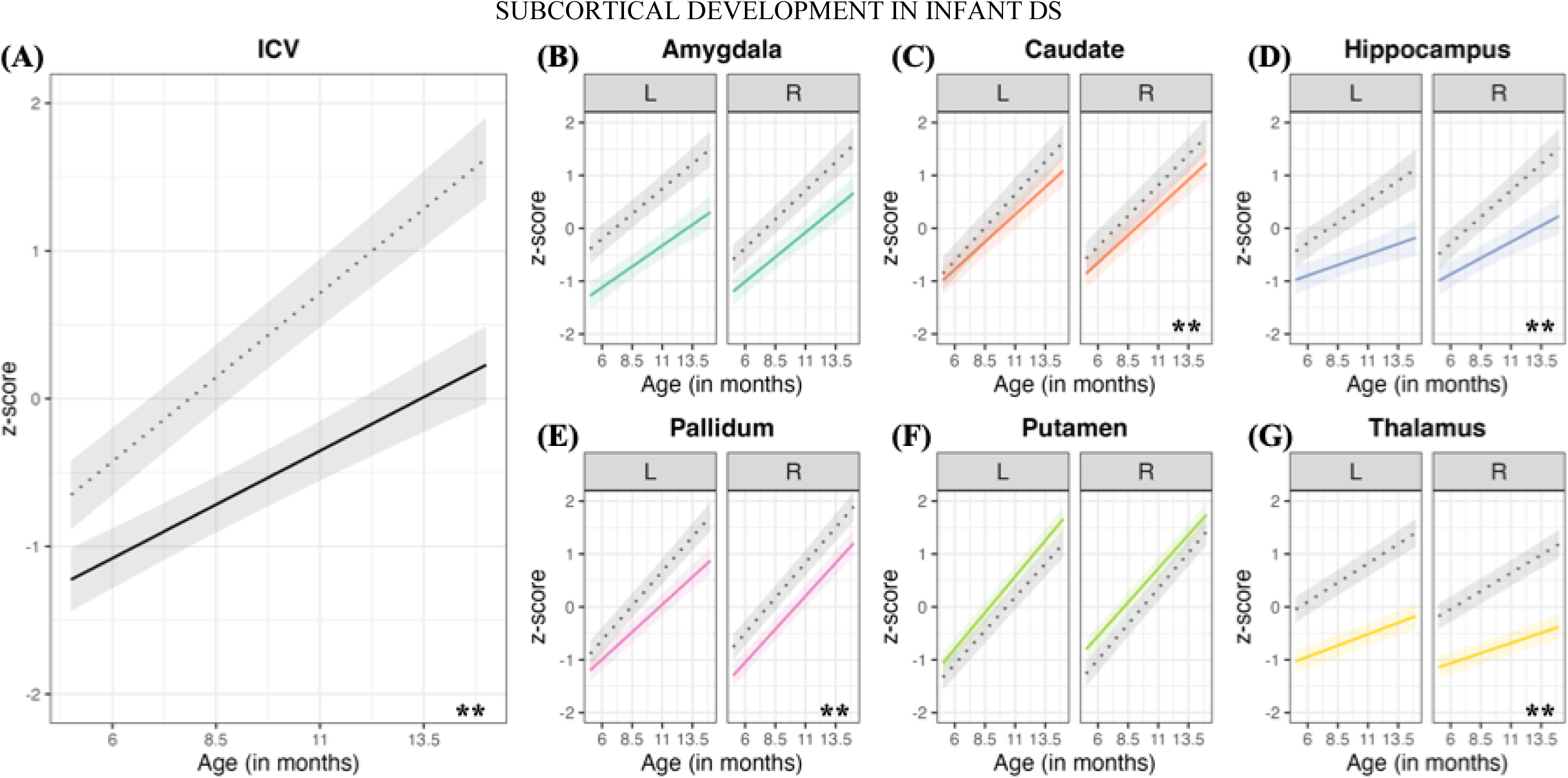
Whole-brain (ICV [black, 2A]) and subcortical regional (amygdala [teal, 2B], caudate [orange, 2C], hippocampus [purple, 2D], pallidum [pink, 2E], putamen [lime green, 2F], thalamus [yellow, 2G]) gray matter development from 6 to 12 months. Predicted changes in gray matter volume (y-axis) across early development (x-axis) in controls (dotted line) and DS (solid line) participants. Asterisks represent significant age ✕ group interactions, indicating growth rate group differences. *ICV = intracranial volume, L = left hemisphere, R = right hemisphere, *** q < .001, ** q < .01, * q < .05* .

### 3.4 Examination of Effects by Age and Hemisphere in Hierarchical Linear Models

To unpack group differences at each time point, the standardized main effects of cohort (DS versus control) for ICV and subcortical volumes were calculated for each time point and right and left hemispheres (**Supplemental Tables 1-3**). In **Figure 3**, standardized effect sizes with confidence intervals display group differences for left and right volumes at 6 and 12 months. The hatched line at 0 allows comparisons of group differences for absolute subcortical volumes.

**Figure 3.**
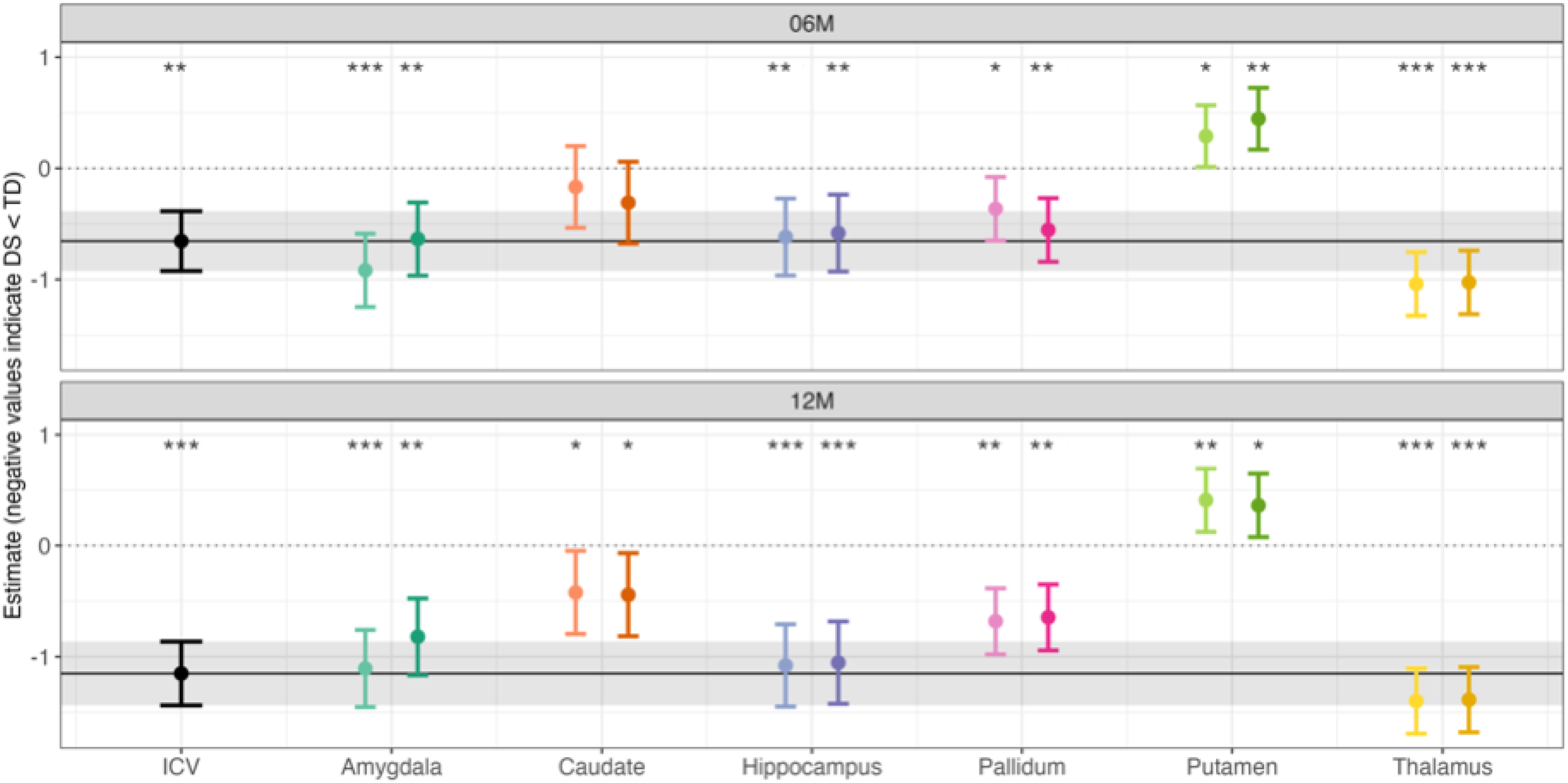
Whole-brain (ICV [black]) and subcortical regional (amygdala [teal], caudate [orange], hippocampus [purple], pallidum [pink], putamen [lime green], thalamus [yellow]) gray matter shown cross-sectionally per visit, controlling for sex and gestational age. Covariate-adjusted absolute values (gestational age, age, sex) from the HLM were used to qualitatively compare the standardized main effect of cohort for each subcortical region to that of ICV (denoted by confidence interval bar), indicating whether DS-associated regional differences were comparable to, larger than, or smaller than overall ICV differences. Standardized effect sizes for cohort are shown at 6 (top) and 12 (bottom) months. Significant differences between control and DS absolute volumes are indicated by asterisks. For a given brain region, light and dark shading differentiate left and right hemispheres, respectively. *ICV = intracranial volume, L = left hemisphere, R = right hemisphere, 06M = 6 months, 12M = 12 months, *** q < .001, ** q < .01, * q < .05*.

At 6 months, the main effect of cohort was significant for ICV, with a standardized effect size of -0.66, indicating a DS-associated reduction in ICV of 0.66 standard deviations below the mean versus control infants (*q*=.002). Significantly lower volumes were also observed for the amygdala, hippocampus, pallidum, and thalamus, with standardized effects from -1.04 for thalamus to -0.36 for pallidum (**Table 3; Supplemental Table 1**). Both right and left caudate volumes were nominally lower in the DS group, but these values did not significantly differ from controls, as the confidence intervals overlapped zero. Left and right putamen uniquely displayed increased volume, with standardized effect sizes of 0.29 (*q*=.047) and 0.45 (*q*=.002), respectively.

Similar profiles across volumes were observed at 12 months (**Supplemental Tables 2-3**), with numerically larger group differences between infants with and without DS. Lower ICV in the DS group showed a standardized effect size of -1.15 (*q*=.001), and more negative effect sizes were observed for amygdala, hippocampus, and pallidum, ranging from -1.40 for left thalamus to -0.42 for left caudate. In contrast to the 6-month time point, 12-month reductions in the left (*q*=.03) and right caudate were significant (*q*=.02). Putamen volumes remained larger in the DS group with standardized effects of 0.41 for left putamen (*q*=.006) and 0.36 for right putamen (*q*=.014). Laterality differences were observed only for the amygdala and did not survive multiple-comparisons correction (*q* = .12).

### 3.5 Qualitative Comparison of Subcortical and ICV Group Differences

We also qualitatively compared DS-associated differences in absolute subcortical volumes with reductions in ICV. In **Figure 3**, the dark line indicates the effect size (with 95% confidence intervals) for lower ICV in DS, a standard of comparison for effect sizes comparing infants with DS and controls across subcortical regions. Subcortical effect sizes within the ICV confidence intervals indicate subcortical volume reductions in DS proportional to reductions in ICV, while effect sizes above or below the confidence intervals suggested enlarged or reduced volumes relative to ICV, respectively.

At 6 and 12 months, lower volumes for the bilateral amygdala, hippocampus, pallidum, and thalamus in DS fell within the 95% confidence intervals for ICV, suggesting that these reductions are proportional to reductions in ICV observed in DS. At 12 months, lower caudate volumes in infants with DS represented a relatively smaller reduction than that observed for ICV. Left and right putamen volumes at both ages were enlarged in infants with DS relative to ICV, with a greater degree of enlargement at 12 than at 6 months.

Supplementary analyses examining subcortical differences when covarying for ICV (**Supplementary Tables 4-7**) were overall congruent with the qualitative approach. Subcortical volumes with group differences similar in magnitude to ICV (e.g., amygdala) generally were not significantly different in these models. The putamen was significant for relative enlargement at both time points (*q*s<.05), as well as the caudate at both 6- and 12-months (*q*s<.05).

## 4. Discussion

In this longitudinal brain imaging study of infants with DS, several observations show continuity with previous literature from older ages, including (1) smaller total brain volume in infants with DS and (2) decreased volume in multiple subcortical structures, with the putamen uniquely showing increased volume, aligning with findings in youth through adulthood.^42,62,86^ Our longitudinal analysis extends prior work by showing that both total brain volumes and most, but not all, subcortical regions exhibit slower growth in infants with DS than in controls. These findings highlight that structural brain differences in DS across the lifespan must be interpreted in the context of regional volumetric differences present by infancy and that regional brain-based biomarkers may inform future clinical trials targeting early development in DS.

### 4.1 DS is Associated with Slower Growth in Total Brain Volume in the First Year of Life

We observed reduced ICV at 6 and 12 months in infants with DS, both when examining raw differences in absolute volumes (13-16% reduction) and when accounting for nested observations and covariates using HLM (−0.66 standardized effect size). These findings correspond to lower total brain volumes reported in the limited infant DS literature.^37,38,42^ Further, both the increased effect size at 12 versus 6 months and the age-by-cohort interactions in HLM support slower ICV growth in DS during the first year of life. Slowed brain growth during a stage of accelerated brain expansion is consistent with early behavioral delays in DS and may contribute to eventual consolidation of delays into intellectual disability.

Existing neuropathological studies suggest that lower ICV and reduced growth may reflect sequelae of cellular alterations observed in postmortem fetal and perinatal studies of DS, such as fewer neurons, shorter dendrites, and impaired cortical layering.^87^ Lower ICV and slower growth could also reflect delayed brain maturation and reduced synaptogenesis during the first year of life, as supported by prior diffusion findings from IBIS-DS, which suggested widespread reduction in neurite density,^88^ as well as morphometric studies of DS indicating slower and less organized cortical development in infancy and childhood.^88,89^ At older ages, lower ICV in children with DS generally shows larger effect sizes^42^ than reported here, suggesting an increasingly divergent neurodevelopmental profile over time. Disambiguating whether alterations in brain morphology reflect delayed brain growth, sequelae of earlier disruptions, or ongoing perturbations related to maturation or degeneration is essential to inform intervention targets for future clinical trials of neurobiologically based treatments for T21.

### 4.2 DS is Primarily Associated with Reduced Subcortical Volumes and Growth in Infancy

The amygdala, hippocampus, thalamus, and pallidum all showed significantly lower absolute volumes at 6 and 12 months in infants with DS, with reductions proportional to the decrease in ICV. DS-associated reduction in absolute amygdala volumes (9.8%-14.8%) aligned with the single available neonatal neuroimaging study of DS,^37^ which found lower absolute amygdala volumes, but not lower volumes relative to overall brain volume. Several studies in children and adolescents with DS also reported smaller amygdala volumes versus controls, including after accounting for reduced total brain volume.^41,59^ In contrast, McCann *et al.* (2021)^42^ noted amygdala enlargement relative to lower total ICV in children with DS between ages 5 and 20 years and an absolute volume increase in a sample of 15 children aged 10-15 years with DS. The agreement of findings in infant studies illustrate the need for developmentally focused, sufficiently powered longitudinal studies to resolve whether inconsistent amygdala findings relate to age or advancing growth trajectories.^41,42,59^

As with the amygdala, smaller absolute hippocampal volumes (9.9-13.2%) were found in infants with DS versus controls, a reduction corresponding with the prior neonatal neuroimaging study in DS^37^ and reduced neurogenesis and neuronal density noted in previous fetal studies.^35,90,91^ As a vital brain region for memory consolidation affected in both DS and Alzheimer’s dementia (a frequent DS comorbidity)^92,93^ the hippocampus has been of great interest in understanding the neurobiology of DS outcomes. Absolute and proportional reductions (relative to total brain) in hippocampal volumes have been observed throughout childhood and adulthood in DS,^41,59–61^ both before and after the onset of Alzheimer’s dementia and its associated hippocampal neuropathology and atrophy.^17,59–61,86^ Taken together, these findings could suggest temporal specificity in the vulnerability of the hippocampus, whereby in infancy, absolute hippocampal volumes, but not whole-brain adjusted volumes, are smaller, followed by later, more substantial reductions measurable via both absolute and relative hippocampal volumes. Such a course could reflect the combined impact of slower early hippocampal growth and subsequent neurodegeneration.

Our study also found large reductions in absolute thalamic volumes (but not in volumes relative to ICV) in infants with DS ranging from 11.8% to 13.1% at 6 and 12 months. This result differs from the relative enlargement previously reported in neonates with DS^37^ and the lack of significant difference in a sample of 16 0-5-year-olds with DS.^42^ However, the observation concurs with smaller thalamus volumes observed when comparing individuals with DS versus controls in studies of 2- to 5-year-olds,^38^ 0- to 18-year-olds,^63^ and 6- to 24-year-olds.^64^ These mixed findings, reminiscent of literature for the amygdala, underscores the need to carefully evaluate the interpretability and generalizability of findings in light of small sample sizes, developmental stage, methodology, and measurement precision. Thalamus volumes, an aggregation of multiple nuclei with different functions, morphological influences, and growth profiles,^63,94^ may also pose unique measurement challenges. More work is needed in DS to establish morphological and volumetric differences in the thalamus and its nuclei, particularly given the thalamus’s role as a relay station for motor, sensory, and limbic functions, all of which are affected in DS.^94^

In addition to reduced volumes, infants with DS also displayed slower growth rates from 6-12 months in the hippocampus, thalamus, and pallidum, suggesting that volumetric differences between groups may widen beyond the first year of life. The hippocampus showed the slowest growth rates in infants with DS (on par with ICV), while the amygdala’s growth rate was numerically but not significantly reduced relative to controls. Notably, the hippocampus is also the structure most affected by Alzheimer’s-related neurodegeneration in DS,^86^ raising the possibility that its vulnerability to the effects of T21 extends across the full lifespan. The presence of regional variation in growth across these structures aligns with studies of typical subcortical development, including faster maturation of the amygdala versus other subcortical structures,^53^ as well as neuropathological studies showing differences in the regional impact of T21 on brain cellularity.^84,95^ Primary analyses of absolute volumes, rather than ICV-adjusted volumes, may also have optimized sensitivity to detect subtler regional influences of T21, while avoiding confounds from possible nonlinear relationships between subcortical volumes and ICV. Incorporating absolute volumes in future research could thus enhance identification of regional brain differences that inform biologically based treatment targets.

### 4.3 Basal Ganglia Structures Show Dissociated Volumetric Differences in Infants with DS

Within the basal ganglia, lower absolute volumes were consistently observed only in the pallidum (6.9%-9.9% at 6 and 12 months), and these differences were less than those for the amygdala, thalamus, and hippocampus. The putamen showed uniquely enlarged absolute volumes in infants with DS (1.5%-5.3%), contrasting with reduced ICV, and was the only subcortical structure to exhibit numerically greater 6-12-month volumetric growth in infants with versus without DS. For the caudate, absolute volumes adjusted for covariates in the HLM were lower at 12 but not 6 months in infants with DS, a shift attributable to slower caudate growth rates in DS. However, because the 12-month reduction in caudate volume was less than that for ICV (**Figure 3**), the caudate was relatively enlarged at 12 months, an observation supported by models controlling for ICV (**Supplementary Table 4-7)**. Although not uniformly observed in DS,^37,41^ caudate enlargement relative to ICV has been reported within all age-stratified subgroups of a sample of 0-22-year-olds with DS.^42^

Referencing volumetric differences for the pallidum and putamen to the DS literature is challenged by historical joint examination of these regions as components of the lentiform nucleus. Within pediatric studies of DS, the lentiform nucleus has generally been enlarged compared to controls,^37,38,41,42,60^ but since only two such studies examined the putamen and pallidum separately,^38,42^ this consensus may not reflect unique properties of each structure. The larger absolute volumes, specifically in the putamen, as well as the slower growth rate for the pallidum in infants with DS, imply that the previous findings for lentiform nucleus volumes in DS may be largely attributable to the putamen. Overall, basal ganglia structures were less likely to show reduced volumes in infants with DS than other subcortical structures, in keeping with prior literature.^42,60^

Finally, relative or absolute enlargements in brain volumes in DS, although often more similar to typical values, may not necessarily reflect compensation or preservation of function. For example, in a study of age-matched adolescents with and without DS,^68^ adolescents with DS exhibited greater gray matter density in the caudate, potentially indicative of less organized brain structure. In infants with fragile X syndrome, enlarged caudate volumes have been quantitatively associated with greater repetitive and restrictive behavior.^43^ In autism, greater putamen volume has also been associated with greater restrictive and repetitive behaviors.^44,96–98^ Putamen enlargement has been proposed to result from dopaminergic dysfunction,^99,100^ which has been observed in DS fetal tissue and animal models.^101,102^ Given the basal ganglia’s role in motor control and the significant motor impairment in DS, future research should examine associations between basal ganglia structure and circuitry and cognitive, motor, and adaptive outcomes in children with DS.^14,103^

### 4.4 Atypical Laterality is Observed for Subcortical Structures During Early Infancy in DS

At 6 months, we observed lateralization differences in infants with DS relative to controls for the amygdala, caudate, pallidum, and putamen, although only the amygdala exhibited differences at 12 months. Previous studies in older children with DS have noted atypical laterality in brain dynamics measured by resting-state EEG,^104^ and several behavioral features of DS are consistent with disrupted functional lateralization, such as atypical handedness,^105,106^ reduced verbal-motor integration,^107^ and reduced efficiency of interhemispheric communication.^108^ Further research is needed to investigate whether these differences provide a developmentally sensitive marker of altered neurodevelopment in DS and relate to functional outcomes.

### 4.5 Implications of Brain Differences for Guiding DS Treatment

The well-defined genetic basis of DS poses unique opportunities for targeted interventions, and considerable progress is being made towards clinical trial readiness. This includes not only the identification of therapeutic targets but also the development of reliable biomarkers for treatment stratification, risk assessment, and monitoring of intervention response. Our work confirms that early subcortical measurements can capture change and regionally specific variation, supporting the feasibility of developmental neuroimaging as a source of non-invasive, objective markers that are sensitive to future treatments. Indeed, mouse models of DS have provided compelling evidence that drugs targeting the Sonic Hedgehog Pathway in early life can impact neurogenesis, connectivity, and function in the hippocampus.^109,110^ Similarly, dysregulated mammalian target of rapamycin (mTOR) signaling pathways^111^ and downregulation of canonical Wnt signaling,^112,113^ due to the overexpression of DYRK1A housed on human chromosome 21,^114^ are implicated in neural development, synaptic formation, and hippocampal plasticity, as well as neuropathology in Alzheimer’s dementia,^115–117^ highlighting therapeutic opportunities across the lifespan. Testing of potential treatments targeting these pathways is underway, including in young children with neurodevelopmental disorders.^118^ This underscores the need for developmentally sensitive markers in childhood and ongoing neurodevelopmental imaging to inform both targeting and monitoring of intervention responses.

### 4.6 Strengths, Limitations, and Future Directions

Our study had numerous strengths, including leveraging a unique, large longitudinal sample of infants with and without DS examined during a period of rapid brain development. Previous DS neuroimaging studies have often relied on retrospective analysis of clinical scans, which may be biased by elevated clinical severity. Our prospective design with standardized data collection thus enhances rigor and generalizability, while the relatively large sample for DS enhances representation of heterogeneity.

To sensitively measure developmental differences in brain volumes, we implemented a new segmentation framework embedding longitudinal processing into the image segmentation step. This advanced technique contrasts with manual segmentations,^40,41,61,66,119,120^ which can introduce bias and error, constraining reproducibility. Further, by using atlases appropriate for a well-defined infant age range, our approach minimized the likelihood of measurement errors from applying atlases from older ages, e.g., tissue mislabeling, registration errors, and distortion due to early, rapid, nonlinear brain growth,^121–123^ and promoted interpretability of findings.

One limitation is that analyses were confined to two time points, rather than estimating trajectories, which could reveal nonlinear changes in rates of brain development over time. The 12-month sample was also smaller than the 6-month sample, a common challenge in longitudinal studies, primarily due to attrition and greater difficulty with unsedated sleeping scans in older infants. However, the demographic characteristics of the 12-month subsample were consistent with those of the full 6-month sample (**Supplementary Table 8**). The use of HLM, which accommodates unbalanced data across time points, also mitigates potential bias due to differential participation. Further, while our sample size is large for DS research, it was not sufficiently powered to test the impact of medical comorbidities that could be associated with brain volumetric differences.^37,42^ Future studies, including ongoing data collection in this study for ages 6-24 months, can expand on our findings via larger sample sizes, additional time points, and examination of brain-behavior associations to probe the impact of aberrant subcortical development in DS on clinical outcomes.

### 4.7 Conclusion

From 6 to 12 months of age, infants with DS exhibited a marked reduction in overall brain size relative to age-matched control infants and slower growth in total brain volume. Varying group differences in subcortical volumes and growth rates support regional specificity in the neural impact of T21 and widening brain volumetric differences in DS during infancy. These results highlight the important role of developmental neuroimaging in characterizing brain-based biomarkers and windows of rapid neurodevelopment for high-impact interventions.

