## SupplementalMaterials for "Variation in infant subcortical brain development from 6 to 12 months in Down syndrome"

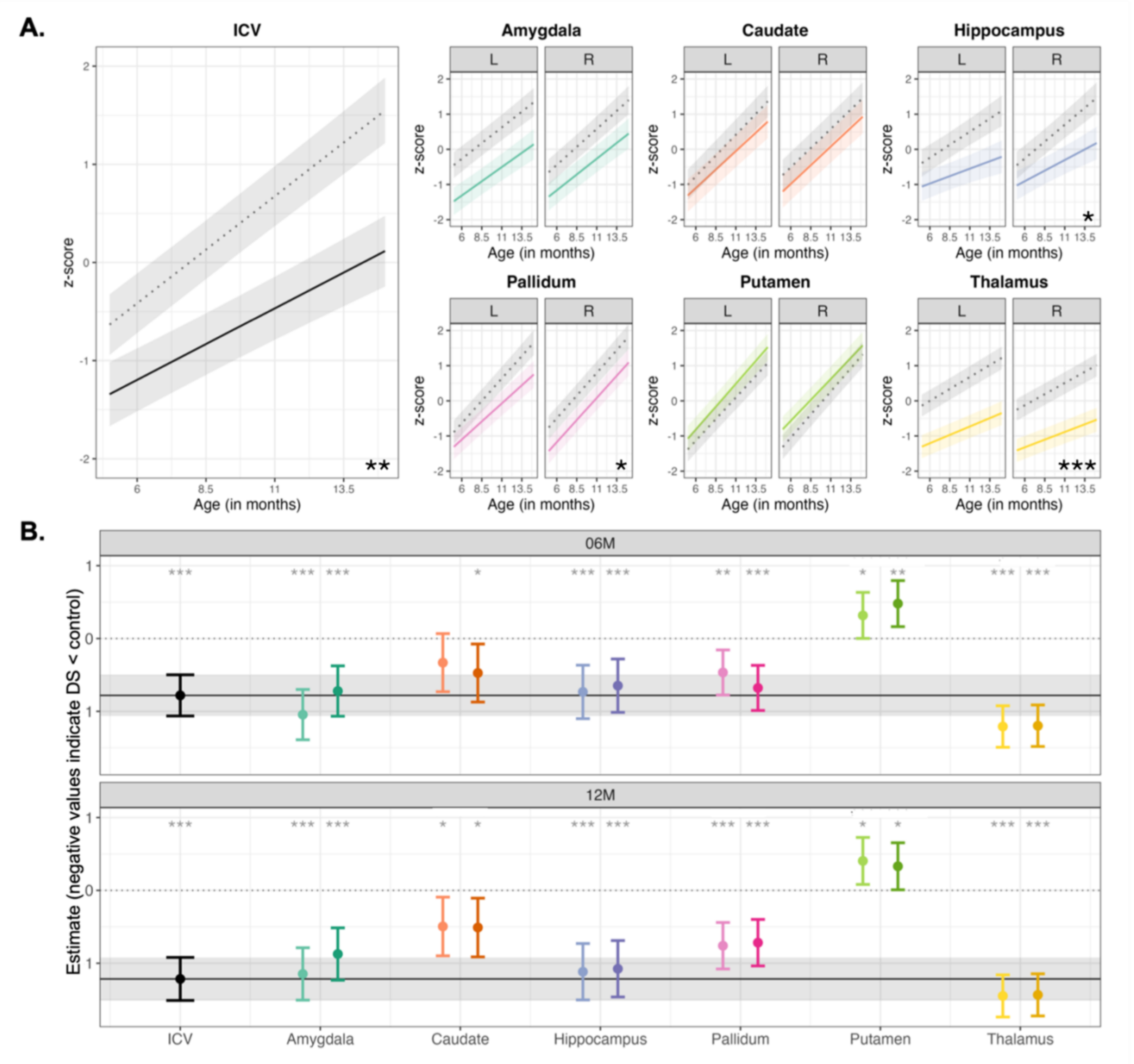


**Supplementary Figure 1.** Unharmonized whole-brain (ICV [black]) and subcortical regional (amygdala [teal], caudate [orange], hippocampus [purple], pallidum [pink], putamen [lime green], thalamus [yellow]) gray matter development from 6 to 12 months. **(A)** Predicted changes in gray matter volume (y-axis) across early development (x-axis) in controls (dotted line) and DS (solid line) participants. **(B)** Standardized effect sizes for cohort at 6 (top) and 12 (bottom) months. Significant differences between control and DS volumes, evaluated with respect to 95% confidence intervals around effect size estimates, are indicated by gray asterisks. For a given brain region, light and dark shading differentiate left and right hemispheres, respectively. Asterisks represent significant age $\times$ group interactions, indicating growth rate group differences in unharmonized results. Harmonized and unharmonized results were nearly identical, indicative of the robust imaging protocol and segmentation approach. *ICV = intracranial volume, L = left hemisphere, R = right hemisphere, 06M = 6 months, 12M = 12 months, *** q < .001, ** q < .01, * q < .05*

| | ICV | | | Amygdala | | | Caudate | | | Hippocampus | | | Pallidum | | | Putamen | | | Thalamus | | |
|---|---|---|---|---|---|---|---|---|---|---|---|---|---|---|---|---|---|---|---|---|---|
| *Predictors* | *Est.* | *SE* | *p* | *Est.* | *SE* | *p* | *Est.* | *SE* | *p* | *Est.* | *SE* | *p* | *Est.* | *SE* | *p* | *Est.* | *SE* | *p* | *Est.* | *SE* | *p* |
| (Intercept) | -0.22 | 0.12 | 0.076 | -0.23 | 0.15 | 0.117 | -0.34 | 0.17 | **0.050** | -0.09 | 0.15 | 0.566 | -0.40 | 0.13 | **0.002** | -0.83 | 0.13 | **<0.001** | 0.15 | 0.13 | 0.240 |
| Sex [Male] | 0.59 | 0.12 | **<0.001**[b] | 0.43 | 0.14 | **0.003**[b] | 0.09 | 0.17 | 0.612 | 0.48 | 0.14 | **0.001**[b] | 0.31 | 0.13 | **0.014**[c] | 0.35 | 0.13 | **0.006**[b] | 0.51 | 0.13 | **<0.001**[b] |
| Gestational age | 0.14 | 0.05 | **0.003**[c] | 0.10 | 0.05 | 0.076 | 0.01 | 0.06 | 0.925 | 0.14 | 0.05 | **0.013**[c] | 0.07 | 0.05 | 0.177 | 0.11 | 0.05 | **0.018**[c] | 0.13 | 0.05 | **0.008**[c] |
| Cohort [DS vs Control] | -0.66 | 0.14 | **<0.001**[b] | -0.64 | 0.17 | **<0.001**[b] | -0.31 | 0.19 | 0.096 | -0.58 | 0.17 | **0.001**[b] | -0.55 | 0.14 | **<0.001**[b] | 0.45 | 0.14 | **0.002**[b] | -1.03 | 0.14 | **<0.001**[a] |
| age (6mo) | 0.23 | 0.01 | **<0.001**[a] | 0.22 | 0.02 | **<0.001**[a] | 0.23 | 0.01 | **<0.001**[a] | 0.20 | 0.02 | **<0.001**[a] | 0.27 | 0.01 | **<0.001**[a] | 0.27 | 0.01 | **<0.001**[a] | 0.14 | 0.01 | **<0.001**[a] |
| Cohort [DS] × age (6mo) | -0.08 | 0.02 | **<0.001**[b] | -0.03 | 0.02 | 0.178 | -0.02 | 0.01 | 0.114 | -0.08 | 0.03 | **0.005**[c] | -0.02 | 0.02 | 0.359 | -0.01 | 0.01 | 0.300 | -0.06 | 0.01 | **<0.001**[b] |
| hemi [L vs R] | | | | 0.17 | 0.08 | **0.029**[c] | -0.27 | 0.05 | **<0.001**[b] | 0.01 | 0.10 | 0.904 | -0.13 | 0.06 | **0.018**[c] | -0.09 | 0.05 | 0.062 | 0.14 | 0.05 | **0.003**[b] |
| Cohort [DS] × hemi [L] | | | | -0.28 | 0.11 | **0.012**[c] | 0.14 | 0.07 | **0.035** | -0.03 | 0.14 | 0.800 | 0.19 | 0.08 | **0.019**[c] | -0.16 | 0.06 | **0.017**[c] | -0.01 | 0.07 | 0.833 |
| age (6mo) × hemi [L] | | | | -0.03 | 0.02 | 0.168 | 0.02 | 0.01 | 0.120 | -0.04 | 0.02 | 0.070 | -0.01 | 0.01 | 0.714 | -0.02 | 0.01 | 0.175 | 0.01 | 0.01 | 0.451 |
| (Cohort [DS] × age (6mo) × hemi [L] | | | | -0.00 | 0.03 | 0.984 | -0.02 | 0.02 | 0.224 | 0.00 | 0.03 | 0.963 | -0.04 | 0.02 | 0.062 | 0.03 | 0.02 | **0.037** | 0.00 | 0.02 | 0.989 |
| **Random Effects** | | | | | | | | | | | | | | | | | | | | | |
| $\sigma^2$ | 0.06 | | | 0.10 | | | 0.03 | | | 0.15 | | | 0.05 | | | 0.03 | | | 0.03 | | |
| $\tau_{00}$ | 0.24 $_{IDs}$ | | | 0.36 $_{IDs}$ | | | 0.56 $_{IDs}$ | | | 0.35 $_{IDs}$ | | | 0.30 $_{IDs}$ | | | 0.30 $_{IDs}$ | | | 0.32 $_{IDs}$ | | |
| ICC | 0.81 | | | 0.79 | | | 0.94 | | | 0.70 | | | 0.86 | | | 0.90 | | | 0.91 | | |
| N | 83 $_{IDs}$ | | | 83 $_{IDs}$ | | | 83 $_{IDs}$ | | | 83 $_{IDs}$ | | | 83 $_{IDs}$ | | | 83 $_{IDs}$ | | | 83 $_{IDs}$ | | |
| Observations | 116 | | | 232 | | | 232 | | | 232 | | | 232 | | | 232 | | | 232 | | |
| Marginal $R^2$ / Conditional $R^2$ | 0.720 / 0.947 | | | 0.584 / 0.912 | | | 0.444 / 0.967 | | | 0.526 / 0.860 | | | 0.656 / 0.951 | | | 0.668 / 0.967 | | | 0.674 / 0.969 | | |

**Supplementary Table 1.** Estimates (Est), standard errors (SE), and p-values (*p*) from hierarchical linear models characterizing early (6-to-12-month) gray matter development in control infants and infants with DS examined at the whole-brain level (ICV) and in six subcortical regions (amygdala, caudate, hippocampus, pallidum, putamen, and thalamus). This table is in addition to Table 3 from the main text, using the left hemisphere as a reference, centered at the 6-month time point. *ICV = intracranial volume, DS = Down syndrome, Hemi = hemisphere.* [a]*q*<.001, [b]*q* <.01, [c]*q* <.05

| | ICV | | | Amygdala | | | Caudate | | | Hippocampus | | | Pallidum | | | Putamen | | | Thalamus | | |
|---|---|---|---|---|---|---|---|---|---|---|---|---|---|---|---|---|---|---|---|---|---|
| *Predictors* | *Est.* | *SE* | *p* | *Est.* | *SE* | *p* | *Est.* | *SE* | *p* | *Est.* | *SE* | *p* | *Est.* | *SE* | *p* | *Est.* | *SE* | *p* | *Est.* | *SE* | *p* |
| (Intercept) | 1.15 | 0.13 | **<0.001** | 1.07 | 0.15 | **<0.001** | 1.05 | 0.17 | **<0.001** | 1.12 | 0.16 | **<0.001** | 1.20 | 0.13 | **<0.001** | 0.79 | 0.13 | **<0.001** | 0.97 | 0.13 | **<0.001** |
| Sex [Male] | 0.59 | 0.12 | **<0.001[a]** | 0.43 | 0.14 | **0.003[b]** | 0.09 | 0.17 | 0.612 | 0.48 | 0.14 | **0.001[b]** | 0.31 | 0.13 | **0.014[c]** | 0.35 | 0.13 | **0.006[b]** | 0.51 | 0.13 | **<0.001[b]** |
| Gestational age | 0.14 | 0.05 | **0.003[c]** | 0.10 | 0.05 | 0.076 | 0.01 | 0.06 | 0.925 | 0.14 | 0.05 | **0.013[c]** | 0.07 | 0.05 | 0.177 | 0.11 | 0.05 | **0.018[c]** | 0.13 | 0.05 | **0.008[c]** |
| Cohort [DS vs Control] | -1.15 | 0.14 | **<0.001[a]** | -0.82 | 0.17 | **<0.001[b]** | -0.44 | 0.19 | **0.020[c]** | -1.05 | 0.19 | **<0.001[a]** | -0.65 | 0.15 | **<0.001[b]** | 0.36 | 0.14 | **0.012[c]** | -1.39 | 0.15 | **<0.001[a]** |
| age (12mo) | 0.23 | 0.01 | **<0.001[a]** | 0.22 | 0.02 | **<0.001[a]** | 0.23 | 0.01 | **<0.001[a]** | 0.20 | 0.02 | **<0.001[a]** | 0.27 | 0.01 | **<0.001[a]** | 0.27 | 0.01 | **<0.001[a]** | 0.14 | 0.01 | **<0.001[a]** |
| Cohort [DS] × age (12mo) | -0.08 | 0.02 | **<0.001[b]** | -0.03 | 0.02 | 0.178 | -0.02 | 0.01 | 0.114 | -0.08 | 0.03 | **0.005[c]** | -0.02 | 0.02 | 0.359 | -0.01 | 0.01 | 0.300 | -0.06 | 0.01 | **<0.001[b]** |
| hemi [L vs R] | | | | 0.01 | 0.09 | 0.929 | -0.16 | 0.05 | **0.004[b]** | -0.25 | 0.11 | **0.020[c]** | -0.17 | 0.06 | **0.010[c]** | -0.18 | 0.05 | **0.001[b]** | 0.19 | 0.05 | **<0.001[b]** |
| Cohort [DS] × hemi [L] | | | | -0.29 | 0.12 | **0.020** | 0.02 | 0.07 | 0.781 | -0.03 | 0.15 | 0.866 | -0.04 | 0.09 | 0.690 | 0.05 | 0.07 | 0.512 | -0.01 | 0.07 | 0.862 |
| age (12mo) × hemi [L] | | | | -0.03 | 0.02 | 0.168 | 0.02 | 0.01 | 0.120 | -0.04 | 0.02 | 0.070 | -0.01 | 0.01 | 0.714 | -0.02 | 0.01 | 0.175 | 0.01 | 0.01 | 0.451 |
| (Cohort [DS] × age (12mo) × hemi [L] | | | | -0.00 | 0.03 | 0.984 | -0.02 | 0.02 | 0.224 | 0.00 | 0.03 | 0.963 | -0.04 | 0.02 | 0.062 | 0.03 | 0.02 | **0.037** | 0.00 | 0.02 | 0.989 |
| **Random Effects** | | | | | | | | | | | | | | | | | | | | | |
| $\sigma^2$ | 0.06 | | | 0.10 | | | 0.03 | | | 0.15 | | | 0.05 | | | 0.03 | | | 0.03 | | |
| $\tau_{00}$ | $0.24_{IDs}$ | | | $0.36_{IDs}$ | | | $0.56_{IDs}$ | | | $0.35_{IDs}$ | | | $0.30_{IDs}$ | | | $0.30_{IDs}$ | | | $0.32_{IDs}$ | | |
| ICC | 0.81 | | | 0.79 | | | 0.94 | | | 0.70 | | | 0.86 | | | 0.90 | | | 0.91 | | |
| N | $83_{IDs}$ | | | $83_{IDs}$ | | | $83_{IDs}$ | | | $83_{IDs}$ | | | $83_{IDs}$ | | | $83_{IDs}$ | | | $83_{IDs}$ | | |
| Observations | 116 | | | 232 | | | 232 | | | 232 | | | 232 | | | 232 | | | 232 | | |
| Marginal $R^2$ / Conditional $R^2$ | 0.720 / 0.947 | | | 0.584 / 0.912 | | | 0.444 / 0.967 | | | 0.526 / 0.860 | | | 0.656 / 0.951 | | | 0.668 / 0.967 | | | 0.674 / 0.969 | | |

**Supplementary Table 2.** Estimates (Est), standard errors (SE), and p-values (*p*) from hierarchical linear models characterizing early (6-to-12-month) gray matter development in control infants and infants with DS examined at the whole-brain level (ICV) and in six subcortical regions (amygdala, caudate, hippocampus, pallidum, putamen, and thalamus). This table is in addition to Table 3 from the main text, using the left hemisphere as a reference, centered at the 12-month time point. *ICV = intracranial volume, DS = Down syndrome, Hemi = hemisphere.* [a]*q*<.001, [b]*q* <.01, [c]*q* <.05

| | ICV | | | Amygdala | | | Caudate | | | Hippocampus | | | Pallidum | | | Putamen | | | Thalamus | | |
|---|---|---|---|---|---|---|---|---|---|---|---|---|---|---|---|---|---|---|---|---|---|
| *Predictors* | *Est.* | *SE* | *p* | *Est.* | *SE* | *p* | *Est.* | *SE* | *p* | *Est.* | *SE* | *p* | *Est.* | *SE* | *p* | *Est.* | *SE* | *p* | *Est.* | *SE* | *p* |
| (Intercept) | 1.15 | 0.13 | **<0.001** | 1.08 | 0.15 | **<0.001** | 0.90 | 0.17 | **<0.001** | 0.86 | 0.16 | **<0.001** | 1.03 | 0.13 | **<0.001** | 0.61 | 0.13 | **<0.001** | 1.16 | 0.13 | **<0.001** |
| Sex [Male] | 0.59 | 0.12 | **<0.001**[b] | 0.43 | 0.14 | **0.003**[b] | 0.09 | 0.17 | 0.612 | 0.48 | 0.14 | **0.001**[b] | 0.31 | 0.13 | **0.014**[b] | 0.35 | 0.13 | **0.006**[b] | 0.51 | 0.13 | **<0.001**[b] |
| Gestational age | 0.14 | 0.05 | **0.003**[c] | 0.10 | 0.05 | 0.076 | 0.01 | 0.06 | 0.925 | 0.14 | 0.05 | **0.013**[c] | 0.07 | 0.05 | 0.177 | 0.11 | 0.05 | **0.018**[c] | 0.13 | 0.05 | **0.008**[c] |
| Cohort [DS vs Control] | -1.15 | 0.14 | **<0.001**[a] | -1.11 | 0.17 | **<0.001**[a] | -0.42 | 0.19 | **0.026**[c] | -1.08 | 0.19 | **<0.001**[a] | -0.68 | 0.15 | **<0.001**[b] | 0.41 | 0.14 | **0.005**[b] | -1.40 | 0.15 | **<0.001**[a] |
| age (12mo) | 0.23 | 0.01 | **<0.001**[a] | 0.19 | 0.02 | **<0.001**[a] | 0.25 | 0.01 | **<0.001**[a] | 0.16 | 0.02 | **<0.001**[a] | 0.26 | 0.01 | **<0.001**[a] | 0.25 | 0.01 | **<0.001**[a] | 0.15 | 0.01 | **<0.001**[a] |
| Cohort [DS] × age (12mo) | -0.08 | 0.02 | **<0.001**[b] | -0.03 | 0.02 | 0.170 | -0.04 | 0.01 | **0.003**[b] | -0.08 | 0.03 | **0.006**[b] | -0.05 | 0.02 | **0.002**[b] | 0.02 | 0.01 | 0.142 | -0.06 | 0.01 | **<0.001**[b] |
| hemi [R vs L] | | | | -0.01 | 0.09 | 0.929 | 0.16 | 0.05 | **0.004**[b] | 0.25 | 0.11 | **0.020**[c] | 0.17 | 0.06 | **0.010**[c] | 0.18 | 0.05 | **0.001**[b] | -0.19 | 0.05 | **<0.001**[b] |
| Cohort [DS] × hemi [R] | | | | 0.29 | 0.12 | **0.020** | -0.02 | 0.07 | 0.781 | 0.03 | 0.15 | 0.866 | 0.04 | 0.09 | 0.690 | -0.05 | 0.07 | 0.512 | 0.01 | 0.07 | 0.862 |
| age (12mo) × hemi [R] | | | | 0.03 | 0.02 | 0.168 | -0.02 | 0.01 | 0.120 | 0.04 | 0.02 | 0.070 | 0.01 | 0.01 | 0.714 | 0.02 | 0.01 | 0.175 | -0.01 | 0.01 | 0.451 |
| (Cohort [DS] × age (12mo) × hemi [R] | | | | 0.00 | 0.03 | 0.984 | 0.02 | 0.02 | 0.224 | -0.00 | 0.03 | 0.963 | 0.04 | 0.02 | 0.062 | -0.03 | 0.02 | **0.037** | -0.00 | 0.02 | 0.989 |
| **Random Effects** | | | | | | | | | | | | | | | | | | | | | |
| $\sigma^2$ | 0.06 | | | 0.10 | | | 0.03 | | | 0.15 | | | 0.05 | | | 0.03 | | | 0.03 | | |
| $\tau_{00}$ | 0.24 $_{IDs}$ | | | 0.36 $_{IDs}$ | | | 0.56 $_{IDs}$ | | | 0.35 $_{IDs}$ | | | 0.30 $_{IDs}$ | | | 0.30 $_{IDs}$ | | | 0.32 $_{IDs}$ | | |
| ICC | 0.81 | | | 0.79 | | | 0.94 | | | 0.70 | | | 0.86 | | | 0.90 | | | 0.91 | | |
| N | 83 $_{IDs}$ | | | 83 $_{IDs}$ | | | 83 $_{IDs}$ | | | 83 $_{IDs}$ | | | 83 $_{IDs}$ | | | 83 $_{IDs}$ | | | 83 $_{IDs}$ | | |
| Observations | 116 | | | 232 | | | 232 | | | 232 | | | 232 | | | 232 | | | 232 | | |
| Marginal $R^2$ / Conditional $R^2$ | 0.720 / 0.947 | | | 0.584 / 0.912 | | | 0.444 / 0.967 | | | 0.526 / 0.860 | | | 0.656 / 0.951 | | | 0.668 / 0.967 | | | 0.674 / 0.969 | | |

**Supplementary Table 3.** Estimates (Est), standard errors (SE), and p-values (*p*) from hierarchical linear models characterizing early (6-to-12-month) gray matter development in control infants and infants with DS examined at the whole-brain level (ICV) and in six subcortical regions (amygdala, caudate, hippocampus, pallidum, putamen, and thalamus). This table is in addition to Table 3 from the main text, using the right hemisphere as a reference, centered at the 12-month time point. *ICV = intracranial volume, DS = Down syndrome, Hemi = hemisphere*. [a]*q*<.001, [b]*q* <.01, [c]*q* <.05

| | ICV | | | Amygdala | | | Caudate | | | Hippocampus | | | Pallidum | | | Putamen | | | Thalamus | | |
|---|---|---|---|---|---|---|---|---|---|---|---|---|---|---|---|---|---|---|---|---|---|
| *Predictors* | *Est.* | *SE* | *p* | *Est.* | *SE* | *p* | *Est.* | *SE* | *p* | *Est.* | *SE* | *p* | *Est.* | *SE* | *p* | *Est.* | *SE* | *p* | *Est.* | *SE* | *p* |
| (Intercept) | 0.22 | 0.12 | 0.076 | 0.10 | 0.11 | 0.365 | -0.44 | 0.13 | **0.001** | 0.03 | 0.14 | 0.824 | -0.41 | 0.11 | **<0.001** | -0.81 | 0.11 | **<0.001** | 0.40 | 0.11 | **<0.001** |
| Sex [Male] | 0.59 | 0.12 | **<0.001**[b] | -0.01 | 0.12 | 0.934 | -0.35 | 0.13 | **0.008**[c] | 0.18 | 0.14 | 0.198 | -0.01 | 0.12 | 0.904 | 0.07 | 0.11 | 0.528 | 0.20 | 0.11 | 0.082 |
| Gestational age | 0.14 | 0.05 | **0.003**[c] | -0.01 | 0.04 | 0.872 | -0.10 | 0.05 | **0.045** | 0.07 | 0.05 | 0.197 | -0.01 | 0.04 | 0.772 | 0.05 | 0.04 | 0.253 | 0.06 | 0.04 | 0.159 |
| Cohort [DS vs Control] | -0.66 | 0.14 | **<0.001**[b] | -0.44 | 0.14 | **0.002**[b] | 0.32 | 0.15 | **0.029**[c] | -0.30 | 0.18 | 0.089 | 0.00 | 0.13 | 0.995 | 0.60 | 0.13 | **<0.001**[b] | -0.69 | 0.13 | **<0.001**[b] |
| age (6mo) | 0.23 | 0.01 | **<0.001**[a] | 0.02 | 0.02 | 0.438 | 0.08 | 0.02 | **<0.001**[a] | 0.04 | 0.03 | 0.150 | 0.13 | 0.02 | **<0.001**[a] | 0.14 | 0.02 | **<0.001**[a] | 0.02 | 0.02 | 0.198 |
| Cohort [DS] × age (6mo) | -0.08 | 0.02 | **<0.001**[b] | 0.03 | 0.02 | 0.124 | 0.02 | 0.01 | 0.089 | -0.03 | 0.03 | 0.235 | -0.01 | 0.02 | 0.746 | 0.06 | 0.01 | **<0.001**[b] | -0.01 | 0.01 | 0.328 |
| ICV z | | | | 0.74 | 0.08 | **<0.001**[a] | 0.75 | 0.07 | **<0.001**[a] | 0.49 | 0.10 | **<0.001**[a] | 0.56 | 0.07 | **<0.001**[a] | 0.47 | 0.07 | **<0.001**[a] | 0.53 | 0.07 | **<0.001**[a] |
| hemi [R vs L] | | | | -0.17 | 0.07 | **0.022**[c] | 0.27 | 0.04 | **<0.001**[c] | -0.01 | 0.10 | 0.903 | 0.13 | 0.05 | **0.011**[c] | 0.09 | 0.04 | **0.048** | -0.14 | 0.04 | **0.002**[c] |
| Cohort [DS] × hemi [R] | | | | 0.28 | 0.11 | **0.009**[c] | -0.14 | 0.06 | **0.016**[c] | 0.03 | 0.14 | 0.798 | -0.19 | 0.08 | **0.012**[c] | 0.16 | 0.06 | **0.011**[c] | 0.01 | 0.06 | 0.822 |
| age (6mo) × hemi [R] | | | | 0.03 | 0.02 | 0.149 | -0.02 | 0.01 | 0.074 | 0.04 | 0.02 | 0.067 | 0.01 | 0.01 | 0.695 | 0.02 | 0.01 | 0.149 | -0.01 | 0.01 | 0.421 |
| (Cohort [DS] × age (6mo) × hemi [R] | | | | 0.00 | 0.03 | 0.984 | 0.02 | 0.01 | 0.163 | -0.00 | 0.03 | 0.963 | 0.04 | 0.02 | **0.045** | -0.03 | 0.02 | **0.027** | -0.00 | 0.02 | 0.988 |
| **Random Effects** | | | | | | | | | | | | | | | | | | | | | |
| $\sigma^2$ | 0.06 | | | 0.09 | | | 0.03 | | | 0.14 | | | 0.04 | | | 0.03 | | | 0.03 | | |
| $\tau_{00}$ | 0.24 IDs | | | 0.19 IDs | | | 0.30 IDs | | | 0.27 IDs | | | 0.21 IDs | | | 0.22 IDs | | | 0.21 IDs | | |
| ICC | 0.81 | | | 0.68 | | | 0.92 | | | 0.65 | | | 0.83 | | | 0.88 | | | 0.88 | | |
| N | 83 IDs | | | 83 IDs | | | 83 IDs | | | 83 IDs | | | 83 IDs | | | 83 IDs | | | 83 IDs | | |
| Observations | 116 | | | 232 | | | 232 | | | 232 | | | 232 | | | 232 | | | 232 | | |
| Marginal/ Conditional $R^2$ | 0.720 / 0.947 | | | 0.736 / 0.915 | | | 0.647 / 0.971 | | | 0.592 / 0.859 | | | 0.744 / 0.955 | | | 0.744 / 0.970 | | | 0.770 / 0.971 | | |

**Supplementary Table 4.** Estimates (Est), standard errors (SE), and p-values (*p*) from hierarchical linear models (HLM) characterizing early (6-to-12-month) gray matter development in infants with Down syndrome and controls, including whole-brain level (ICV) and six subcortical regions (amygdala, caudate, hippocampus, pallidum, putamen, and thalamus). ICV is included as a covariate in these secondary analyses, for which the right hemisphere as the reference hemisphere and centering is at the 6-month time point. In this HLM, the putamen and caudate show relative enlargement in DS, while the thalamus is relatively smaller in DS. For the amygdala, hippocampus, and pallidum, adjusting for ICV attenuates significant group effects observed with absolute volumes in primary analyses. This is expected in cases for which subcortical group differences are proportional to group differences in ICV and relationships between subcortical regions and ICV show consistent scaling across age and group. *ICV = intracranial volume, DS = Down syndrome, Hemi = hemisphere*. [a]$q<.001$, [b]$q<.01$, [c]$q<.05$

| | ICV | | | Amygdala | | | Caudate | | | Hippocampus | | | Pallidum | | | Putamen | | | Thalamus | | |
|---|---|---|---|---|---|---|---|---|---|---|---|---|---|---|---|---|---|---|---|---|---|
| *Predictors* | *Est.* | *SE* | *p* | *Est.* | *SE* | *p* | *Est.* | *SE* | *p* | *Est.* | *SE* | *p* | *Est.* | *SE* | *p* | *Est.* | *SE* | *p* | *Est.* | *SE* | *p* |
| (Intercept) | -0.22 | 0.12 | 0.076 | -0.07 | 0.11 | 0.552 | -0.18 | 0.13 | 0.167 | 0.02 | 0.14 | 0.888 | -0.28 | 0.11 | **0.013** | -0.73 | 0.11 | **<0.001** | 0.27 | 0.11 | **0.014** |
| Sex [Male] | 0.59 | 0.12 | **<0.001[b]** | -0.01 | 0.12 | 0.934 | -0.35 | 0.13 | **0.008[c]** | 0.18 | 0.14 | 0.198 | -0.01 | 0.12 | 0.904 | 0.07 | 0.11 | 0.528 | 0.20 | 0.11 | 0.082 |
| Gestational age | 0.14 | 0.05 | **0.003[c]** | -0.01 | 0.04 | 0.872 | -0.10 | 0.05 | **0.045** | 0.07 | 0.05 | 0.197 | -0.01 | 0.04 | 0.772 | 0.05 | 0.04 | 0.253 | 0.06 | 0.04 | 0.159 |
| Cohort [DS vs Control] | -0.66 | 0.14 | **<0.001[b]** | -0.16 | 0.14 | 0.260 | 0.18 | 0.15 | 0.225 | -0.26 | 0.18 | 0.133 | -0.19 | 0.13 | 0.153 | 0.75 | 0.13 | **<0.001[a]** | -0.68 | 0.13 | **<0.001[a]** |
| age (6mo) | 0.23 | 0.01 | **<0.001[a]** | 0.05 | 0.02 | 0.057 | 0.06 | 0.02 | **0.001[a]** | 0.09 | 0.03 | **0.004[b]** | 0.14 | 0.02 | **<0.001[a]** | 0.16 | 0.02 | **<0.001[a]** | 0.01 | 0.02 | 0.428 |
| Cohort [DS] × age (6mo) | -0.08 | 0.02 | **<0.001[b]** | 0.04 | 0.02 | 0.119 | 0.04 | 0.01 | **0.002[b]** | -0.04 | 0.03 | 0.214 | 0.03 | 0.02 | 0.055 | 0.03 | 0.01 | 0.061 | -0.01 | 0.01 | 0.320 |
| ICV z | | | | 0.74 | 0.08 | **<0.001[a]** | 0.75 | 0.07 | **<0.001[a]** | 0.49 | 0.10 | **<0.001[a]** | 0.56 | 0.07 | **<0.001[a]** | 0.47 | 0.07 | **<0.001[a]** | 0.53 | 0.07 | **<0.001[a]** |
| hemi [L vs R] | | | | 0.17 | 0.07 | **0.022[c]** | -0.27 | 0.04 | **<0.001[b]** | 0.01 | 0.10 | 0.903 | -0.13 | 0.05 | **0.011[c]** | -0.09 | 0.04 | **0.048** | 0.14 | 0.04 | **0.002[b]** |
| Cohort [DS] × hemi [L] | | | | -0.28 | 0.11 | **0.009[c]** | 0.14 | 0.06 | **0.016[c]** | -0.03 | 0.14 | 0.798 | 0.19 | 0.08 | **0.012[c]** | -0.16 | 0.06 | **0.011[c]** | -0.01 | 0.06 | 0.822 |
| age (6mo) × hemi [L] | | | | -0.03 | 0.02 | 0.149 | 0.02 | 0.01 | 0.074 | -0.04 | 0.02 | 0.067 | -0.01 | 0.01 | 0.695 | -0.02 | 0.01 | 0.149 | 0.01 | 0.01 | 0.421 |
| (Cohort [DS] × age (6mo) × hemi [L] | | | | -0.00 | 0.03 | 0.984 | -0.02 | 0.01 | 0.163 | 0.00 | 0.03 | 0.963 | -0.04 | 0.02 | **0.045** | 0.03 | 0.02 | **0.027** | 0.00 | 0.02 | 0.988 |
| **Random Effects** | | | | | | | | | | | | | | | | | | | | | |
| $\sigma^2$ | 0.06 | | | 0.09 | | | 0.03 | | | 0.14 | | | 0.04 | | | 0.03 | | | 0.03 | | |
| $\tau_{00}$ | 0.24 $_{IDs}$ | | | 0.19 $_{IDs}$ | | | 0.30 $_{IDs}$ | | | 0.27 $_{IDs}$ | | | 0.21 $_{IDs}$ | | | 0.22 $_{IDs}$ | | | 0.21 $_{IDs}$ | | |

| | | | | | | | |
|---|---|---|---|---|---|---|---|
| ICC | 0.81 | 0.68 | 0.92 | 0.65 | 0.83 | 0.88 | 0.88 |
| N | 83 IDs | 83 IDs | 83 IDs | 83 IDs | 83 IDs | 83 IDs | 83 IDs |
| Observations | 116 | 232 | 232 | 232 | 232 | 232 | 232 |
| Marginal $R^2$ / Conditional $R^2$ | 0.720 / 0.947 | 0.736 / 0.915 | 0.647 / 0.971 | 0.592 / 0.859 | 0.744 / 0.955 | 0.744 / 0.970 | 0.770 / 0.971 |

**Supplementary Table 5.** Estimates (Est), standard errors (SE), and p-values (*p*) from hierarchical linear models (HLM) characterizing early (6-to-12-month) gray matter development in infants with Down syndrome and controls, including whole-brain level (ICV) and six subcortical regions (amygdala, caudate, hippocampus, pallidum, putamen, and thalamus). ICV is included as a covariate in these secondary analyses, for which the left hemisphere is the reference hemisphere and centering is at the 6-month time point. In this HLM, the putamen shows relative enlargement in DS, while the thalamus is relatively smaller in DS. For the amygdala, caudate, hippocampus, and pallidum, adjusting for ICV attenuates significant group effects observed with absolute volumes in primary analyses. This is expected in cases for which subcortical group differences are proportional to group differences in ICV and relationships between subcortical regions and ICV show consistent scaling across age and group. *ICV = intracranial volume, DS = Down syndrome, Hemi = hemisphere*. [a]*q*<.001, [b]*q* <.01, [c]*q* <.05

| | ICV | | | Amygdala | | | Caudate | | | Hippocampus | | | Pallidum | | | Putamen | | | Thalamus | | |
|---|---|---|---|---|---|---|---|---|---|---|---|---|---|---|---|---|---|---|---|---|---|
| *Predictors* | *Est.* | *SE* | *p* | *Est.* | *SE* | *p* | *Est.* | *SE* | *p* | *Est.* | *SE* | *p* | *Est.* | *SE* | *p* | *Est.* | *SE* | *p* | *Est.* | *SE* | *p* |
| (Intercept) | 1.15 | 0.13 | **<0.001** | 0.22 | 0.15 | 0.161 | 0.02 | 0.15 | 0.877 | 0.29 | 0.19 | 0.125 | 0.38 | 0.14 | **0.008** | 0.06 | 0.14 | 0.678 | 0.54 | 0.13 | **<0.001** |
| Sex [Male] | 0.59 | 0.12 | **<0.001**[a] | -0.01 | 0.12 | 0.934 | -0.35 | 0.13 | **0.008**[c] | 0.18 | 0.14 | 0.198 | -0.01 | 0.12 | 0.904 | 0.07 | 0.11 | 0.528 | 0.20 | 0.11 | 0.082 |
| Gestational age | 0.14 | 0.05 | **0.003**[c] | -0.01 | 0.04 | 0.872 | -0.10 | 0.05 | **0.045** | 0.07 | 0.05 | 0.197 | -0.01 | 0.04 | 0.772 | 0.05 | 0.04 | 0.253 | 0.06 | 0.04 | 0.159 |
| Cohort [DS vs Control] | -1.15 | 0.14 | **<0.001**[b] | -0.23 | 0.17 | 0.171 | 0.46 | 0.16 | **0.006**[c] | -0.50 | 0.21 | **0.018**[c] | -0.03 | 0.16 | 0.839 | 0.96 | 0.15 | **<0.001**[a] | -0.78 | 0.15 | **<0.001**[b] |
| age (12mo) | 0.23 | 0.01 | **<0.001**[a] | 0.02 | 0.02 | 0.438 | 0.08 | 0.02 | **<0.001**[a] | 0.04 | 0.03 | 0.150 | 0.13 | 0.02 | **<0.001**[a] | 0.14 | 0.02 | **<0.001**[a] | 0.02 | 0.02 | 0.198 |
| Cohort [DS] × age (12mo) | -0.08 | 0.02 | **<0.001**[b] | 0.03 | 0.02 | 0.124 | 0.02 | 0.01 | 0.089 | -0.03 | 0.03 | 0.235 | -0.01 | 0.02 | 0.746 | 0.06 | 0.01 | **<0.001**[b] | -0.01 | 0.01 | 0.328 |
| ICV z | | | | 0.74 | 0.08 | **<0.001**[a] | 0.75 | 0.07 | **<0.001**[a] | 0.49 | 0.10 | **<0.001**[a] | 0.56 | 0.07 | **<0.001**[a] | 0.47 | 0.07 | **<0.001**[a] | 0.53 | 0.07 | **<0.001**[a] |
| hemi [R vs L] | | | | -0.01 | 0.08 | 0.926 | 0.16 | 0.05 | **0.001**[b] | 0.25 | 0.11 | **0.019**[c] | 0.17 | 0.06 | **0.006**[b] | 0.18 | 0.05 | **<0.001**[b] | -0.19 | 0.05 | **<0.001**[b] |
| Cohort [DS] × hemi [R] | | | | 0.29 | 0.12 | **0.015** | -0.02 | 0.06 | 0.750 | 0.03 | 0.15 | 0.865 | 0.04 | 0.08 | 0.670 | -0.05 | 0.07 | 0.486 | 0.01 | 0.07 | 0.853 |
| age (12mo) × hemi [R] | | | | 0.03 | 0.02 | 0.149 | -0.02 | 0.01 | 0.074 | 0.04 | 0.02 | 0.067 | 0.01 | 0.01 | 0.695 | 0.02 | 0.01 | 0.149 | -0.01 | 0.01 | 0.421 |
| (Cohort [DS] × age (12mo) × hemi [R] | | | | 0.00 | 0.03 | 0.984 | 0.02 | 0.01 | 0.163 | -0.00 | 0.03 | 0.963 | 0.04 | 0.02 | **0.045** | -0.03 | 0.02 | **0.027** | -0.00 | 0.02 | 0.988 |
| **Random Effects** | | | | | | | | | | | | | | | | | | | | | |
| $\sigma^2$ | 0.06 | | | 0.09 | | | 0.03 | | | 0.14 | | | 0.04 | | | 0.03 | | | 0.03 | | |
| $\tau_{00}$ | 0.24 IDs | | | 0.19 IDs | | | 0.30 IDs | | | 0.27 IDs | | | 0.21 IDs | | | 0.22 IDs | | | 0.21 IDs | | |
| ICC | 0.81 | | | 0.68 | | | 0.92 | | | 0.65 | | | 0.83 | | | 0.88 | | | 0.88 | | |
| N | 83 IDs | | | 83 IDs | | | 83 IDs | | | 83 IDs | | | 83 IDs | | | 83 IDs | | | 83 IDs | | |
| Observations | 116 | | | 232 | | | 232 | | | 232 | | | 232 | | | 232 | | | 232 | | |
| Marginal $R^2$ / Conditional $R^2$ | 0.720 / 0.947 | | | 0.736 / 0.915 | | | 0.647 / 0.971 | | | 0.592 / 0.859 | | | 0.744 / 0.955 | | | 0.744 / 0.970 | | | 0.770 / 0.971 | | |

**Supplementary Table 6.** Estimates (Est), standard errors (SE), and p-values (*p*) from hierarchical linear models (HLM) characterizing early (6-to-12-month) gray matter development in infants with Down syndrome and controls, including whole-brain level (ICV) and six subcortical regions (amygdala, caudate, hippocampus, pallidum, putamen, and thalamus). ICV is included as a covariate in these secondary analyses, for which the right hemisphere is the reference hemisphere and centering is at the 12-month time point. In this HLM, the putamen and caudate show relative enlargement in

DS, while the hippocampus and thalamus are relatively smaller in DS. For the amygdala and pallidum, adjusting for ICV attenuates significant group effects observed with absolute volumes in primary analyses. This is expected in cases for which subcortical group differences are proportional to group differences in ICV and relationships between subcortical regions and ICV show consistent scaling across age and group. *ICV = intracranial volume, DS = Down syndrome, Hemi = hemisphere*. [a]$q<.001$, [b]$q<.01$, [c]$q<.05$

| | ICV | | | Amygdala | | | Caudate | | | Hippocampus | | | Pallidum | | | Putamen | | | Thalamus | | |
|---|---|---|---|---|---|---|---|---|---|---|---|---|---|---|---|---|---|---|---|---|---|
| *Predictors* | *Est.* | *SE* | *P* | *Est.* | *SE* | *p* | *Est.* | *SE* | *p* | *Est.* | *SE* | *p* | *Est.* | *SE* | *p* | *Est.* | *SE* | *p* | *Est.* | *SE* | *p* |
| (Intercept) | 1.15 | 0.13 | **<0.001** | 0.21 | 0.15 | 0.177 | 0.18 | 0.15 | 0.240 | 0.55 | 0.19 | **0.004** | 0.55 | 0.14 | **<0.001** | 0.24 | 0.14 | 0.086 | 0.35 | 0.13 | **0.009** |
| Sex [Male] | 0.59 | 0.12 | **<0.001[b]** | -0.01 | 0.12 | 0.934 | -0.35 | 0.13 | **0.008[c]** | 0.18 | 0.14 | 0.198 | -0.01 | 0.12 | 0.904 | 0.07 | 0.11 | 0.528 | 0.20 | 0.11 | 0.082 |
| Gestational age | 0.14 | 0.05 | **0.003[c]** | -0.01 | 0.04 | 0.872 | -0.10 | 0.05 | **0.045** | 0.07 | 0.05 | 0.197 | -0.01 | 0.04 | 0.772 | 0.05 | 0.04 | 0.253 | 0.06 | 0.04 | 0.159 |
| Cohort [DS vs Control] | -1.15 | 0.14 | **<0.001[a]** | 0.05 | 0.17 | 0.763 | 0.44 | 0.16 | **0.008[c]** | -0.48 | 0.21 | **0.025[c]** | 0.00 | 0.16 | 0.982 | 0.91 | 0.15 | **<0.001[b]** | -0.76 | 0.15 | **<0.001[b]** |
| age (12mo) | 0.23 | 0.01 | **<0.001[a]** | 0.05 | 0.02 | 0.057 | 0.06 | 0.02 | **0.001[b]** | 0.09 | 0.03 | **0.004[b]** | 0.14 | 0.02 | **<0.001[b]** | 0.16 | 0.02 | **<0.001[b]** | 0.01 | 0.02 | 0.428 |
| Cohort [DS] × age (12mo) | -0.08 | 0.02 | **<0.001[b]** | 0.04 | 0.02 | 0.119 | 0.04 | 0.01 | **0.002[b]** | -0.04 | 0.03 | 0.214 | 0.03 | 0.02 | 0.055 | 0.03 | 0.01 | 0.061 | -0.01 | 0.01 | 0.320 |
| ICV z | | | | 0.74 | 0.08 | **<0.001[a]** | 0.75 | 0.07 | **<0.001[a]** | 0.49 | 0.10 | **<0.001[b]** | 0.56 | 0.07 | **<0.001[a]** | 0.47 | 0.07 | **<0.001[a]** | 0.53 | 0.07 | **<0.001[a]** |
| hemi [L vs R] | | | | 0.01 | 0.08 | 0.926 | -0.16 | 0.05 | **0.001[b]** | -0.25 | 0.11 | **0.019[c]** | -0.17 | 0.06 | **0.006[b]** | -0.18 | 0.05 | **<0.001[b]** | 0.19 | 0.05 | **<0.001[b]** |
| Cohort [DS] × hemi [L] | | | | -0.29 | 0.12 | **0.015** | 0.02 | 0.06 | 0.750 | -0.03 | 0.15 | 0.865 | -0.04 | 0.08 | 0.670 | 0.05 | 0.07 | 0.486 | -0.01 | 0.07 | 0.853 |
| age (12mo) × hemi [L] | | | | -0.03 | 0.02 | 0.149 | 0.02 | 0.01 | 0.074 | -0.04 | 0.02 | 0.067 | -0.01 | 0.01 | 0.695 | -0.02 | 0.01 | 0.149 | 0.01 | 0.01 | 0.421 |
| (Cohort [DS] × age (12mo) × hemi [L] | | | | -0.00 | 0.03 | 0.984 | -0.02 | 0.01 | 0.163 | 0.00 | 0.03 | 0.963 | -0.04 | 0.02 | **0.045** | 0.03 | 0.02 | **0.027** | 0.00 | 0.02 | 0.988 |
| **Random Effects** | | | | | | | | | | | | | | | | | | | | | |
| $\sigma^2$ | 0.06 | | | 0.09 | | | 0.03 | | | 0.14 | | | 0.04 | | | 0.03 | | | 0.03 | | |
| $\tau_{00}$ | 0.24 $_{IDs}$ | | | 0.19 $_{IDs}$ | | | 0.30 $_{IDs}$ | | | 0.27 $_{IDs}$ | | | 0.21 $_{IDs}$ | | | 0.22 $_{IDs}$ | | | 0.21 $_{IDs}$ | | |
| ICC | 0.81 | | | 0.68 | | | 0.92 | | | 0.65 | | | 0.83 | | | 0.88 | | | 0.88 | | |
| N | 83 $_{IDs}$ | | | 83 $_{IDs}$ | | | 83 $_{IDs}$ | | | 83 $_{IDs}$ | | | 83 $_{IDs}$ | | | 83 $_{IDs}$ | | | 83 $_{IDs}$ | | |
| Observations | 116 | | | 232 | | | 232 | | | 232 | | | 232 | | | 232 | | | 232 | | |
| Marginal $R^2$ / Conditional $R^2$ | 0.720 / 0.947 | | | 0.736 / 0.915 | | | 0.647 / 0.971 | | | 0.592 / 0.859 | | | 0.744 / 0.955 | | | 0.744 / 0.970 | | | 0.770 / 0.971 | | |

**Supplementary Table 7.** Estimates (Est), standard errors (SE), and p-values (*p*) from hierarchical linear models (HLM) characterizing early (6-to-12-month) gray matter development in infants with Down syndrome and controls, including whole-brain level (ICV) and six subcortical regions (amygdala, caudate, hippocampus, pallidum, putamen, and thalamus). ICV is included as a covariate in these secondary analyses, for which the left hemisphere is the reference hemisphere and centering is at the 12-month time point. In this HLM, the putamen and caudate show relative enlargement in DS, while the hippocampus and thalamus are relatively smaller in DS. For the amygdala and pallidum, adjusting for ICV attenuates significant group effects observed with absolute volumes in primary analyses. This is expected in cases for which subcortical group differences are proportional to group differences in ICV and relationships between subcortical regions and ICV show consistent scaling across age and group. *ICV = intracranial volume, DS = Down syndrome, Hemi = hemisphere.* $^{a}q<.001$, $^{b}q<.01$, $^{c}q<.05$

| | DS | | | Control | | |
|---|---|---|---|---|---|---|
| | **6 mo (n=39)** | **12 mo (n=20)** | **6 vs. 12 mo p-value** | **6 mo (n=37)** | **12 mo (n=20)** | **6 vs. 12 mo p-value** |
| **Sex, n (%)** | | | | | | |
| Male | 15 (38.5%) | 7 (35.0%) | 1.000 | 21 (56.8%) | 12 (60.0%) | 1.000 |
| Female | 24 (61.5%) | 13 (65.0%) | | 16 (43.2%) | 8 (40.0%) | |
| **Gestational age (weeks), M (SD)** | 37.99 (1.43) | 37.88 (1.47) | .778 | 39.10 (1.18) | 39.09 (1.26) | .964 |
| **Age at 6-mo scan (months), M (SD)** | 6.88 (1.06) | — | — | 6.50 (0.47) | — | — |
| **Age at 12-mo scan (months), M (SD)** | — | 12.66 (1.24) | — | — | 12.57 (0.64) | — |

**Supplementary Table 8**. Demographic characteristics did not differ between subjects with usable data at 6 months and those at 12 months, indicating the 12-month subsamples are representative of their respective broader 6-month cohorts. Fisher's exact test was used for sex; the independent-samples t-test for gestational age.
